# Spatiotemporal patterns of theta-band activity during rapid-eye movement sleep: a magnetoencephalography analysis

**DOI:** 10.1101/2024.10.01.616032

**Authors:** Vasiliki Provias, Monika Schönauer, Steffan Gais, Phillipe Albouy, Jan Born, Til O. Bergmann, Emily B.J. Coffey

## Abstract

Theta oscillations (4-8 Hz) in frontal cortical regions are present to different degrees across states of consciousness. In sleep, theta is prominent in periods of rapid eye-movement (REM) sleep. Theta has been linked to processes of memory consolidation; however, its mechanistic contribution specifically during REM sleep is not well understood. Interestingly, in the wake state, frontal theta activity increases during effortful cognitive tasks involving executive functions such as working memory, hinting at similarities in circuitry, and potentially, function. The aim of the present work is to create a spatially resolved, whole-brain characterisation of REM oscillatory activity in healthy human subjects, distinguishing theta from neighbouring frequency bands, differentiating substages of REM sleep (phasic and tonic REM), and comparing REM theta to that which is evoked during a working memory task. To that end, we analysed magneto- and electroencephalography (M/EEG) data recorded during overnight sleep in 10 healthy subjects, and similar data from 17 healthy subjects who performed a working memory task, using a novel whole-brain, source-localised MEG approach. Our results show that (i) theta activity has a frontal midline topography that is distinct from those of other prominent frequency bands in REM (delta, alpha, beta), (ii) theta activity in frontal midline regions is best observed within a focused 5-7 Hz range, separating it from occipital alpha activity, (iii) REM theta is dominant over the frontal midline but is also observed in several sub-cortical areas, (iv) theta is more widespread in tonic than phasic REM sleep, and (v) the focused frontal midline theta pattern observed in REM phasic sleep is the most similar of all observed sleep substages to theta evoked by a working memory task. These results enhance our understanding of theta physiology in REM sleep and suggest future targets for research into REM’s role in learning and memory.

## 1 Statement of significance

This study provides insights into the role of theta oscillations in REM sleep and their potential links to memory processing. By using a whole-brain, source-localized approach, the research identifies distinct patterns of theta activity in frontal midline regions during REM sleep and highlights similarities between REM and wakeful working memory tasks. These findings suggest that theta oscillations may play a broader role in memory consolidation across different states of consciousness. Future research should explore the mechanisms underlying these oscillations and their potential relevance to cognitive and memory disorders, ultimately advancing our understanding of the connections between sleep, learning, and memory. This work offers new targets for investigating the role of REM sleep in human brain function using causal methods.

## 2 Introduction

Sleep plays an important role in memory consolidation — the process of transforming initially fragile memory traces into strong and stable representations (Diekelmann & Born, 2010). There is substantial evidence supporting the role of non-rapid-eye movement (NREM) sleep in memory consolidation, but the specific contribution of rapid-eye movement (REM) sleep and the mechanisms by which REM might contribute to memory consolidation is less well-understood (Ackermann & Rasch, 2014; Brodt et al., 2023; Fernandez & Lüthi, 2020; Klinzing et al., 2019). Although experimental methods that allow for causal inference such as optogenetics (e.g., Boyce et al., 2016) and brain stimulation (e.g., Lim et al., 2009) may be necessary to confirm the contribution of specific circuits in REM’s putative consolidation roles, insights into where key circuits might be and what functions they might be involved in can first be gained by exploring REM sleep physiology in the whole brain. However, we currently lack a frequency-resolved whole brain characterisation of neural activity in REM sleep.

EEG studies of REM sleep indicate that brain activity exhibits low-amplitude, mixed-frequency patterns closely resembling wakefulness (Ackermann & Rasch, 2014; Brodt et al., 2023). Several frequency bands are observed to be amplified during REM sleep, including delta (1-4 Hz), theta (4-8 Hz), alpha (8-12 Hz), beta (18-30 Hz), and gamma (30-100 Hz) (Cantero et al., 2003; Cape et al., 2000; Montgomery et al., 2008; Nishida et al., 2009; Scheffzük et al., 2011; Simor et al., 2016, 2020, 2021; Vijayan et al., 2017). Theta oscillations have been of particular interest in recent research (Boyce et al., 2016, 2017; Hammer et al., 2021; Harrington et al., 2021; Vijayan et al., 2017), and have been implicated in memory consolidation (Boyce et al., 2016; Nishida et al., 2009; Popa et al., 2010). In one of the few causal studies on this topic, Boyce et al. (2016) demonstrated that optogenetic suppression of medial septum neurons, which generate and pace hippocampal theta rhythms during REM sleep, impaired memory for object recognition and contextual fear in mice. Human studies have suggested that scalp-recorded REM theta over frontal regions is particularly important for emotional memory consolidation (Nishida et al., 2009). Observations such as these have led researchers to propose that theta rhythms during REM sleep may facilitate the offline interaction and encoding of disparate brain regions, enhancing the consolidation of emotional memories (Goldstein & Walker, 2014; Nishida et al., 2009). A significant challenge in testing these ideas and advancing our understanding of REM’s role in memory is that focused, invasive studies in non-human animals allow for spatial and temporal specificity with a limited field of view, whereas studies looking at complex activities of relevance for human behaviour tend to use very global measures of neural activity that offer little insight into its neural basis.

Greater brain coverage and spatial specificity of neural activity in REM can be revealed by positron emission tomography (PET), which indexes metabolic activity, and blood oxygen level dependent functional magnetic resonance imaging (BOLD fMRI), which indexes regional changes in blood oxygenation. Both techniques yield similar results: a broad network of brain regions show greater activity during REM sleep as compared to NREM sleep. Cortical areas that are particularly active in REM sleep include the medial prefrontal cortex (mPFC), inferior frontal gyrus, anterior cingulate cortex (ACC), superior parietal cortex, precuneus, and unimodal sensory areas, and subcortical regions including the amygdala, entorhinal cortex, paralimbic-limbic areas, thalamus, basal ganglia, and pontine tegmentum (Braun et al., 1997; Maquet et al., 2005; Wehrle et al., 2005, 2007). While PET and fMRI studies provide essential spatial information about brain activity, they do not provide information concerning the frequency-specificity of REM-related activity, which makes it difficult to isolate and differentiate networks that communicate within specific frequency bands. It is unclear from these methods which regions may be generating theta activity and communicating with one another in the theta frequency band.

The neural generators of the behaviourally-relevant theta activity observed in human studies is thus only coarsely understood. Studies using time-resolved methods rarely have sufficient sensor density and coverage to accurately map activity onto the brain to resolve the generators of these signals (with a few exceptions, e.g., Baird et al., 2018; Cantero et al., 2003; Simor et al., 2016). Using EEG and intracranial recordings, REM theta-band activity has been detected in the hippocampus and frontal regions, including the ACC and the dlPFC (Cantero et al., 2003; Simor et al., 2016; Vijayan et al., 2017). In the wake state, theta generation has also been reported in the hippocampus, extra-hippocampal structures such as the septal complex, entorhinal cortex, and pedunculopontine tegmentum, as well as cortical regions like the medial prefrontal cortex (mPFC) (Pignatelli et al., 2012). A full-brain analysis of REM theta using neuroimaging methods that allow for resolution of both spatial and frequency information would favour a better understanding of the brain regions and networks that are specific to the theta frequency band and distinct from adjacent bands such as alpha, which may have an overlapping frequency distribution.

Of relevance to understanding REM’s roles, accruing evidence suggests that it is not a homogeneous state but instead can be divided into ‘phasic’ and ‘tonic’ substages, each exhibiting distinct features (Simor et al., 2016, 2020, 2021). Roughly 20-30% of a REM episode consists of phasic events, characterised by bursts of eye movements, heightened cortical activity, and prominent theta and gamma frequencies on EEG (Aserinsky, 1971; Simor et al., 2020). In contrast, tonic REM is marked by the absence of eye movements, and a greater presence of alpha and beta waves (Simor et al., 2020). Phasic and tonic REM sleep substages may serve distinct functions, with differences in environmental alertness and information processing during these substages (Simor et al., 2020; Takahara et al., 2009; Wehrle et al., 2007). As people are less responsive to external stimuli during phasic REM sleep, it may act as an internally focused processing state, potentially facilitating complex cognitive processes (Simor et al., 2016). Therefore, a whole-brain characterisation of theta in REM sleep should examine these substages individually, and to better understand what is unique about REM sleep, compare them with other finer-grained analyses of NREM sleep which contain neural events such as sleep spindles and slow oscillations that have known relationships to theta band activity (Gonzalez et al., 2018; Schreiner et al., 2018).

Further understanding theta’s role in REM sleep processes could benefit from studying frontal theta activity in humans during wakefulness, based on the idea that neural networks interconnecting particular brain structures and sub-serving related functions are constrained by the biological properties of their neural substrates. Frontal theta activity increases during demanding cognitive tasks and resting states (Albouy et al., 2017; Cantero et al., 2003; Capilla et al., 2022; Maurer et al., 2015; Niso et al., 2016). For example, working memory — the cognitive process responsible for the temporary holding and manipulation of information — has been linked to frontal midline theta oscillations in humans (for a review, see Hsieh and Ranganath, 2014). It is compelling to speculate that prefrontal cortical theta oscillations in working memory and those observed during REM sleep involve related cognitive functions, for example in holding in mind sensory or conceptual information, as might occur in a working memory task, when replaying memories during memory consolidation, and perhaps during dreaming (Chow et al., 2013). While testing such a hypothesis would likely involve a series of dedicated experiments using a wide range of neuroscience techniques and tools, a valuable starting point would be to analyse the similarities between frontal theta patterns in (phasic and tonic) REM sleep and those produced during known cognitive processes, such as working memory.

The aim of the present work is to create a spatially resolved, whole-brain characterisation of REM oscillatory activity in healthy human subjects, distinguishing theta from neighbouring frequency bands, differentiating substages of REM sleep (i.e., phasic and tonic REM), and comparing REM theta to that which is evoked during a working memory task. To that end, we analysed electroencephalography EEG/MEG data recorded during overnight sleep in 10 healthy subjects, and similar data from 17 healthy subjects who performed a working memory task, using a novel whole-brain approach that takes advantage of more accurate source-localisation afforded by magnetoencephalography as compared with electroencephalography (Baillet, 2017). We believe that the results will be useful for advancing our understanding of REM sleep’s functional significance and its role in memory consolidation.

## 3 Methods

### 3.1 Participants

#### 3.1.1 Sleep study

We obtained EEG/MEG recordings during nocturnal sleep from 10 healthy participants who were selected for being comfortable in a supine sleeping position. The subjects’ age range was 20-28 (mean: 25.1, SD: 2.6), and six were female. Subjects reported having no sleep-related disorders, not to have changed time zones in the 6 weeks preceding the experiment, and not to be engaged in shift-work. Chronotype was assessed with the Munich Chronotype Questionnaire to adjust the sleep period to the subject’s habitual sleep rhythm. Participants were asked to get up an hour earlier than usual on experimental days, to refrain from napping, and to abstain from consuming alcohol, caffeine, or other substances affecting central nervous system function. The study was approved by the Ethics commission of the University of Tübingen, and written informed consent was obtained for all subjects.

#### 3.1.2 Working memory study

We obtained a separate dataset of MEG/EEG recordings from participants performing an auditory working memory task (Albouy et al., 2017). The study involved 17 healthy participants, including five females (mean age: 28.12, SD: 3.86 years; range: 21-33 years). All participants reported normal hearing and no history of neurological or psychiatric disorders. They provided written informed consent and received monetary compensation for their participation. Each participant underwent a preliminary session to check for potential MEG artefacts. Ethical approval was obtained from the Ethics Review Board of the Montreal Neurological Institute (NEU-14-043) and the Comité d’Éthique de la Recherche en Arts et en Sciences of Université de Montŕeal (CERAS-2014-15-251-D), and written informed consent was obtained for all subjects.

### General experimental design

#### 3.1.3 Sleep study

Each subject in the sleep dataset slept in the MEG scanner for a total of five nights as part of a learning study with other experimental objectives; one adaptation night, followed immediately by the first experimental night. Three subsequent experimental nights took place with intervals of about a week. For the present analysis, one complete experimental night was analysed for each subject, selected for having relatively continuous sleep, and according to availability at the time of analysis. On experimental nights, subjects came to the MEG centre three hours before their sleep time, changed into comfortable sleeping clothes, and were fitted with electrodes. They performed a learning task involving photographs presented at different screen locations (results not reported herein) followed by a memory retrieval task, while seated in the MEG scanner for approximately an hour (note that because sleep data were selected arbitrarily with respect to the behavioural training, the group-level physiological results reported in the present work are not expected to be greatly dependent upon it). The MEG system was then moved to a supine orientation and furnished with bedding, cushioned pads were placed so as to limit head movement, and the lights were turned off. Subjects were awakened after approximately 6 hours of sleep, counted from the first K-complex (slow oscillation) or sleep spindle observed in the EEG recording.

#### 3.1.4 Working memory study

The working memory study involved Transcranial Magnetic Stimulation (TMS); however, our analyses examined data exclusively from the sham condition (i.e., no active TMS session). Participants performed two auditory melodic discrimination tasks: simple melodies and manipulation melodies. Both of these tasks involved detecting if the pitch of a single tone among a series of 3 tones changed in the second melody (with pitch changes of 2 or 3 semitones). In the simple task, participants compared two melodies to determine if a single tone in the second melody was altered. In the manipulation task, the tones of the second melody were rearranged so that the final tone became the first. We used data from both the simple and manipulation tasks to enhance the robustness and reliability of our analyses by providing a larger number of trials; the difference as regards brain activity lies in stronger activation of the left intra-parietal sulcus, a brain region that is involved specifically in transforming mental representations, in the manipulation condition observed in direct comparision between the conditions; this difference has been characterized in the same dataset and reported (see Albouy et al., 2017 for additional details).

### EEG and MEG measurement

For the sleep study, polysomnography (PSG) and MEG data were simultaneously recorded and synchronised using a CTF MEG system (Omega 275, CTF MEG Neuro Innovations Inc.) and its in-built EEG system. PSG was recorded using 12 electrodes at a sampling frequency of 1171.9 Hz, including EEG at C3 and C4 (10–20 International System), electromyogram (left and right jaw muscle), electrooculogram (horizontal and vertical), and electrocardiogram. A reference electrode was affixed to the nose, and the ground electrode was placed on the collarbone. MEG data were recorded from 270 channels (axial gradiometers). Head position was tracked using three head position indicator coils placed anterior to the ears and in the centre of the forehead. Head shape was digitised using a Polhemus Isotrak (Polhemus Inc., VT, USA) for alignment with the subject’s anatomical T1-weighted MRI. For the auditory working memory study, the same model of CTF MEG system was used, but without PSG recording. The sampling rate was set to 1200 Hz with a filter bandwidth of 0–150 Hz. EEG data, not reported herein, were simultaneously acquired from 62 channels and the positions of the EEG electrodes were estimated using the same digitizer system (Polhemus Isotrack).

### Anatomical basis of distributed source models

For both the sleep and working memory studies, T1-weighted 1 mm^3^ isotropic MRI images were acquired for each participant to maximise the accuracy of source reconstruction. In one subject from the sleep study, the MRI was unavailable, so a standard MNI152 brain was used as a substitute, with the relative location of fiducial points estimated using photographs and head anatomy. MRI scans were automatically segmented using Freesurfer and imported into Brainstorm to serve as the basis for anatomical source modelling (Fischl, 2012).

### Sleep scoring and substage identification

To create a basic reference of sleep marcroarchitecture, polysomnographic data were band-pass filtered (0.1-40 Hz) and scored according to Rechtschaffen and Kales’ criteria (Kales & Rechtschaffen, 1968). Two raters independently scored each night’s sleep, based on the C4 electrode, and a third rater then reviewed the two and reconciled any discrepancies. Dividing human sleep into 30 second visually-scored epochs has an extensive history in sleep research and thus is a useful starting point; however, these epochs are relatively long and may encompass multiple neural events with different sources and functions. As several of our research questions on REM sleep concern distinct substates (phasic, tonic; Simor et al., 2020), and because NREM sleep has distinct events such as sleep spindles and slow oscillations that do have known relationships to theta band activity (Gonzalez et al., 2018; Schreiner et al., 2018), meaning that combining them might obscure similarities and differences to activity present in REM sleep and wakefulness in our analyses, we adopted a microarchitecture approach both for both REM and NREM sleep.

To this end, we performed a finer-grained scoring by selecting 5 s non-overlapping windows that were free of movement artefacts and which best and unambiguously represented the sleep substages of interest. For REM, epochs were identified which either clearly contained eye movements (‘REM phasic’) or no eye movements (‘REM tonic’). Using the original 30 s epochs as a reference for the presence of lighter NREM sleep, we marked 3 types of epochs: those in which a spindle occurred but not a K-complex (‘N2 Spindle’), those in which a K-complex occurred (‘N2 K-complex’), and those in which neither was evident (‘N2 Plain’). Note that we adopt the American Academy of Sleep Medicine ‘N2’ abbreviation in this paper to denote epochs in lighter NREM sleep (Iber, 2007), as it is more recognizable to many researchers and is practically equivalent to Rechtschaffen and Kales’ stage 2 sleep (‘S2’). In 30 s epochs classified as deep NREM sleep, 5 s windows in which clear slow wave activity (i.e. high amplitude, 0.5-2 Hz oscillations) was evident for the full duration were selected (slow wave sleep; ’SWS’). These selected epochs were not continuous sub-classifications of the 30 s sleep stages-classification, but rather represented a smaller subset of the most representative neural activity for each substage. The total number of epochs included by sleep substages in the analysis is reported in Table 1.

**Table 1:**
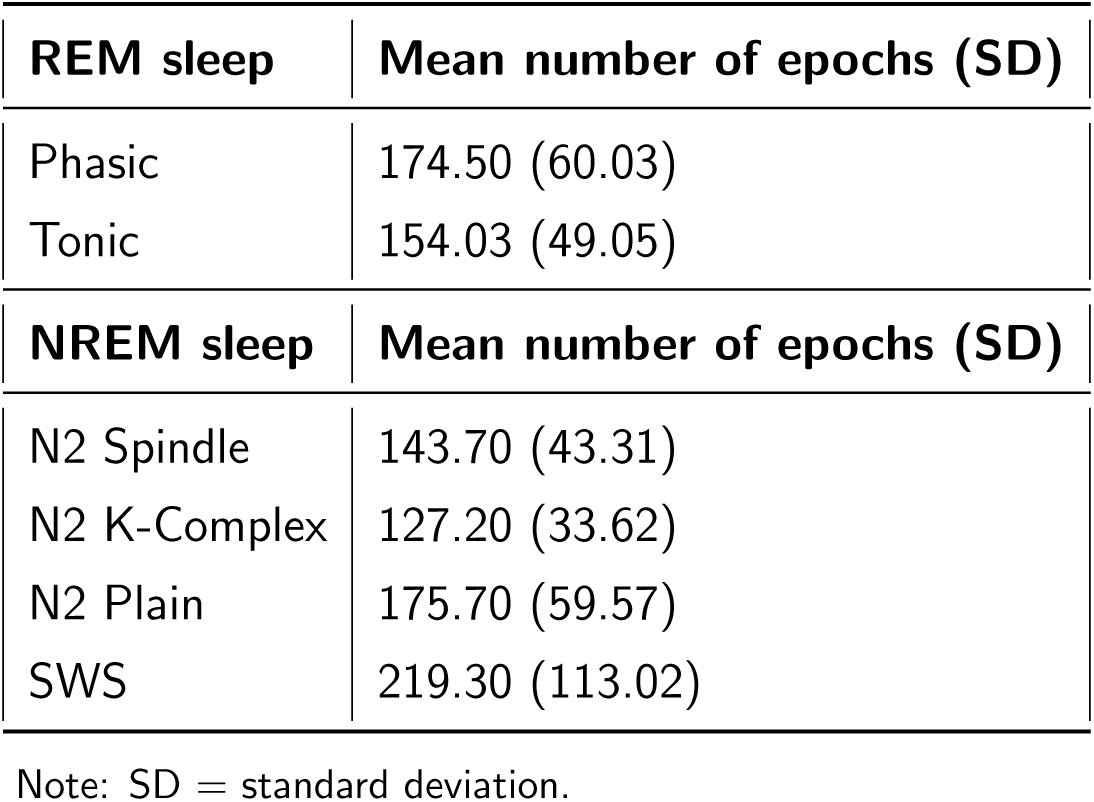
Mean number of 5 s epochs for each sleep stage across all subjects.

| **REM sleep** | **Mean number of epochs (SD)** |
| --- | --- |
| Phasic | 174.50 (60.03) |
| Tonic | 154.03 (49.05) |
| **NREM sleep** | **Mean number of epochs (SD)** |
| N2 Spindle | 143.70 (43.31) |
| N2 K-Complex | 127.20 (33.62) |
| N2 Plain | 175.70 (59.57) |
| SWS | 219.30 (113.02) |

Note: SD = standard deviation.

### MEG sleep analysis

Data analysis was performed with Brainstorm (Tadel et al., 2011) and using custom MATLAB scripts (The Mathworks Inc., MA, USA). Cardiac artefacts were removed using Brainstorm’s in-built source signal space projection algorithm, using the procedure recommended by the software’s authors: projectors were removed when they captured at least 10% of the variance of the signal and the topography matched those of cardiac origin upon visual inspection. Artefacts caused by eye movements were not removed for the sleep data analysis. Blinks and rapid eye movements do not occur in NREM sleep, and because REMs are highly correlated with REM neural activity (Miyauchi et al., 2009), removing co-occurring signals could remove or distort oscillatory activity that is of interest to our research questions. Data were then filtered between 0.1-100 Hz, notch filtered at 50 Hz (power line noise), downsampled to 500 Hz, and epoched based on the sleep substage identidication procedure described above. Using the imported Freesurfer brain segmentation, an overlapping spheres volume head model was computed for each subject based on a regular isotropic 5 mm grid. This forward model explains how an electric current flowing in the brain would be recorded at the level of the sensors, with fair accuracy (Tadel et al., 2011). A noise covariance matrix was computed from 2 min empty room recordings taken before each session for 6 subjects. Recordings were not available for four datasets; we used instead an average computed from the others (these strongly resembled one another as all data were collected on the same instrument during the same time period). The inverse imaging model estimates the distribution of brain currents that accounts for data recorded at the sensors. We computed the MNE source distribution with unconstrained sources and default Brainstorm parameters for each epoch (Gramfort et al., 2014). The MNE source model is simple, robust to noise and model approximations, is frequently used in literature, and has previously been shown them to be sensitive to deep brain sources in the brainstem and thalamus, using a similar analysis pipeline (Coffey et al., 2016, 2021).

### Defining regions of interest

We constructed a whole-brain volume atlas for each subject, comprising 415 regions. The purpose of this step was to keep computational costs reasonable via dimensionality reduction, while ensuring regions of interest (ROIs) were of relatively homogeneous size and shape (i.e., avoiding very large or elongated regions of interest which could reduce signal clarity), and ensuring full brain coverage. Cortical and subcortical regions were derived from the Atlas of Intrinsic Connectivity of Homotopic Areas (AICHA), a functional brain atlas derived from resting-state fMRI data (Joliot et al., 2015). The cerebellum was added using the Automated Anatomical Labelling (AAL) atlas, consisting of 26 regions, including the vermis (Rolls et al., 2020). Additionally, subcortical structures of known relevance to REM sleep were selected a priori on the basis of prior neuroimaging studies described in the introduction, namely the lateral geniculate nuclei (LGN) of the thalamus, the pedunculopontine nucleus (PPN), and the medial septum (Boyce et al., 2016; Gott et al., 2017; Rye, 1997). The LGN was created as a volume scout in standard space based on the FSL’s Harvard–Oxford Cortical and Subcortical Structural probabilistic atlases with coordinates for the left LGN at L = -22, -32, 0 and for the right LGN at R = 24, -30, 0 (Desikan et al., 2006). The PPN and medial septum were defined on the MNI template with reference to gross anatomy, which was subsequently transformed into native space and visually inspected for alignment with individual anatomy. The PPN (L = -4.5, -29.7, -17, R = 4.6, -29.7, -16), each of which (left, right) had a mean volume of 0.096 cm^3^ (SD: 0.107). The medial septum was segmented manually according to Butler et al., (2014) (centred at 0.7, 4.7, 1.9), with a mean volume of 0.552 cm^3^ (SD: 0.126). For each analysis, data were extracted for each of three directions produced by the volume-based MEG model (x, y, z), and subjected to further analysis according to the research question (described below).

### MEG working memory analysis

To compare our sleep data with the topographies of known waking function, we analysed data from 17 healthy young adults who performed an auditory working memory task in which participants had to listen to three tones (encoding period), hold the auditory stimuli in mind (retention period), and finally be prompted via a second set of 3 tones for a match-mismatch decision (retrieval period) (Albouy et al., 2017). In Albouy et al., theta band power was found to be elevated in dorsal and frontal cortical regions during a retention period in which participants held musical tones in mind (as compared to encoding and retrieval periods, in which only temporal auditory regions were active). Dorsal theta activity was higher in trials in which participants subsequently made a correct behavioural choice using the retained information; these findings confirm the behavioural relevance of the frontal theta activity observed in this dataset.

The present analysis was done according to the same steps outlined above for the sleep analysis, with several accommodations; instead of 5 s epochs, we used 3.5 s epochs which was the length of the retention period, and thus the longest interval without the influence of sensory information. This analysis followed the same steps outlined above, with several adjustments: instead of using 5 s epochs, we used 3.5 s epochs, which corresponded to the time interval in which participants retained auditory stimuli in memory before being probed for accuracy (and is of sufficient length nonetheless to characterise the strength of oscillations low theta range, encompassing at least 12 cycles at ∼4 Hz). For the statistical analysis, we created a median topography of working memory for each subject. Each subject’s working memory topography was then compared with a median topography averaged across all sleep dataset subjects, from each sleep stage, using Pearson correlation coefficients. The Pearson correlation coefficient is a straightforward method for assessing the strength and direction of linear relationships between two continuous variables, making it well-suited for our objective of comparing topographical distributions. To assess the significance of similarity between topographies at the group level, we conducted Wilcoxon signed-rank tests, comparing the correlation values to zero, with an alpha level of 0.001.

### Spectral analysis

For spectral analysis, we used an irregular-resampling auto-spectral analysis (IRASA) algorithm (Wen & Liu, 2016). This method can be used to separate fractal (1/f) and oscillatory components in the power spectrum of neurophysiological signals based on their temporal and spectral characteristics, which likely arise from different mechanisms and may differ across sleep stages (Wen & Liu, 2016). We further calculated the percent difference of oscillatory values over fractal values using the following formula:

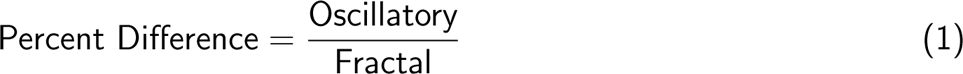

This step normalises oscillatory power for each subject, revealing the less pronounced oscillations typically masked by 1/f noise and reducing the effects of inter-subject variability in EEG/MEG signal strength (Feld et al., 2021). Spectra for each epoch were averaged across the three directions (x, y, z).

### Statistical analysis

To analyse REM versus NREM sleep and phasic versus tonic REM substages, we employed linear mixed-effects modeling (LMEs). This method is suitable for our analysis because it accounts for the nested structure of our data, with includes thousands of epochs for each of a relatively low number of subjects (Schielzeth et al., 2020). LME analyses were conducted in R using the *lmerTest* and *emmeans* packages (Bates et al., 2015; Lenth & Lenth, 2018).

To identify differences in theta activity during REM sleep versus NREM sleep, we conducted an LME analysis at the trial-epoch level, using Condition (i.e., REM and NREM) as a fixed effect and Subjects as a random effect. As we had a large imbalance in the numbers of 5 s epochs by sleep stage (Table 1), we randomly selected an equal number of NREM epochs at the subject level to match those in the REM. The formula used for the analysis was: value ∼ condition (1 + condition | subject). To identify differences in theta activity during phasic and tonic REM substages, we conducted an LME analysis at the trial-epoch level, using conditions (i.e., phasic and tonic) as fixed effects and subjects as a random effect. The formula used for the analysis was: value ∼ condition (1 + condition | subject). LME models were created for each region of interest (ROI), incorporating the condition variable, which depending on the analysis, was either the sleep stage (Figure 3 and 4) or frequency band (Figure 1 reported in Supplementary Materials) as a fixed effect. For each LME model, we visually inspected the histograms and quantile-quantile (Q-Q) plots of the residuals to check for deviations from normality and homoscedasticity. Deviations were addressed by removing outliers, defined as values lying beyond 1.5 times the interquartile range (i.e., below Q1 and above Q3), as in previous work (Jourde et al., 2024). We then conducted an analysis of variance (ANOVA) on each model to assess the significance of the condition variable. The *emmeans* package was used to perform post-hoc comparisons and identify specific regions where significant differences existed between conditions. To control for multiple comparisons, we applied false discovery rate (FDR) correction. Regions showing significant differences (p < 0.05) in oscillatory activity across different conditions were identified and plotted.

**Figure 1:**
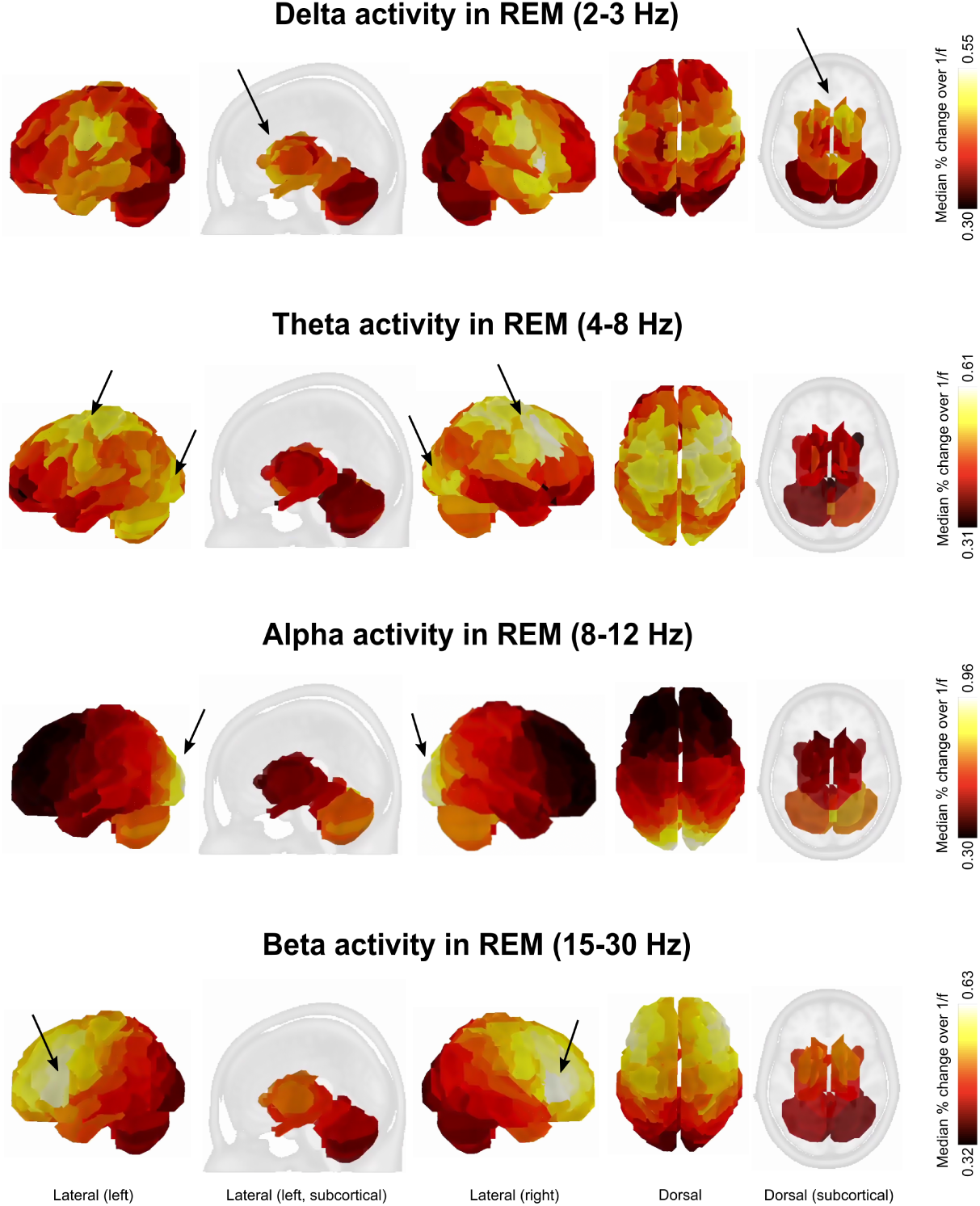
Whole-brain topographies depicting the median oscillatory activity in the delta (2-3 Hz), theta (4-8 Hz), alpha (8-12 Hz), and beta (15-30 Hz) frequency bands during REM sleep, combining both phasic and tonic substages across participants. From left to right, views are left lateral cortical, left lateral subcortical, right lateral cortical, dorsal cortical, and dorsal subcortical. Arrows highlight prominent patterns. Colour scales are scaled within each band to emphasize prominent patterns (ranges shown at right).

### 3.2 Whole-brain topographies

Two types of whole-brain spatial topographical maps were created depending on the research question. First, to observe patterns of oscillatory activity in the whole brain across participants for different frequency bands, we created spatial topographies of the median percent change of oscillatory over fractal activity across all subjects. The colour scale for the median maps was determined based on the minimum and maximum values within the frequency bands of interest for the specific sleep stage being analysed. The working memory topography was also based on this scale. The maximum and minimum values used for all median topographies can be found in Table 1 in Supplementary Materials, and are indicated on colour bars for the main images. Different amplitude ranges have been selected for each frequency band to improve visibility of topographic patterns, noting that a direct comparison of oscillatory amplitude across frequency bands (Figure 1) or sleep stages (Figure 5) is not an intended aim of this research (and would have to be addressed using a different approach). Second, to address questions concerning differences between sleep stages and substages, we created whole-brain topographies of significant F-statistic values obtained from LME models onto the brain. Statistical significance was determined through FDR-corrected p-values, with a significance level (alpha) set at 0.05. The colour scale for these topographies was based on the range of significant F-statistic values (noted on figures). Brain regions that did not reach significance are depicted in grey.

## 4 Results

### 4.1 Sleep scoring

Based on Rechtschaffen and Kales’ sleep scoring criteria (30 s epochs), subjects spent an average of 152.4 min in sleep stage 2 (SD: 14.1), 84.5 min in stages 3 and 4 combined (SD: 17.2), and 58.4 min in REM (SD: 21.0), indicating that participants were able to sleep successfully in the MEG environment. The mean and standard deviation of the number of 5 s epochs selected as most representative of each sleep substage are presented in Table 1. Only these data were used in the analyses.

### 4.2 Defining a frequency range for studying theta activity in REM sleep

#### 4.2.1 Topographies of prominent frequency bands in REM sleep

Figure 1 depicts the median oscillatory activity across subjects in the (commonly-used) 4-8 Hz theta frequency band, as well as delta (2-3 Hz), alpha (8-12 Hz), and beta (15-30 Hz) bands for comparative purposes, during REM sleep and combining both phasic and tonic substages across participants. Visual inspection reveals that beta activity is predominant bilaterally in the lateral frontal regions (in line with previous intracranial work; Vijayan et al., 2017), while alpha is localized to occipital regions (in line with expectations for alpha activity in wakefulness as in Niso et al., 2016). Delta shows both cortical and subcortical activity, potentially suggesting a qualitative difference with the adjacent theta band. In the theta band, two groupings of high oscillatory activity are observed: one in frontal and dorsal regions and another in the posterior regions, encompassing the occipital lobe and cerebellum. Notably, the activity in the posterior regions of the 4-8 Hz theta band resembles the alpha band (8-12 Hz) pattern.

#### 4.2.2 Separating frontal theta and occipital alpha network in REM sleep

The relationship between the topographic patterns in the theta-to-alpha range can be better appreciated via the finer-grained topographies presented in Figure 2A, and in the spectra extracted from sample regions of interest from the frontal and occpital activity peaks depicted in 2B. The amplitude of oscillatory activity within the occipital pattern has a sharp peak around 8.5 Hz, as is expected for the alpha range, whereas in the frontal midline area it has a broader peak at about 5-7 Hz. The topography averaged across the commonly-used 4-8 Hz theta range includes contributions from both of these patterns, with greater influence of the occipital lobe towards the upper upper end of the range (for a direct statistical comparison of brain regions preferentially exhibiting theta versus alpha band activity during REM sleep within the 4-8 Hz range, please refer to Figure 1 in Supplementary Materials). The commonly-used 4-8 Hz range appears to capture two areas of activity with different spectral profiles. To reduce the overlap with alpha activity, which has a known occipital origin (Niso et al., 2016)), we focus on a narrower range of 5-7 Hz in subsequent analyses concerning theta.

**Figure 2:**
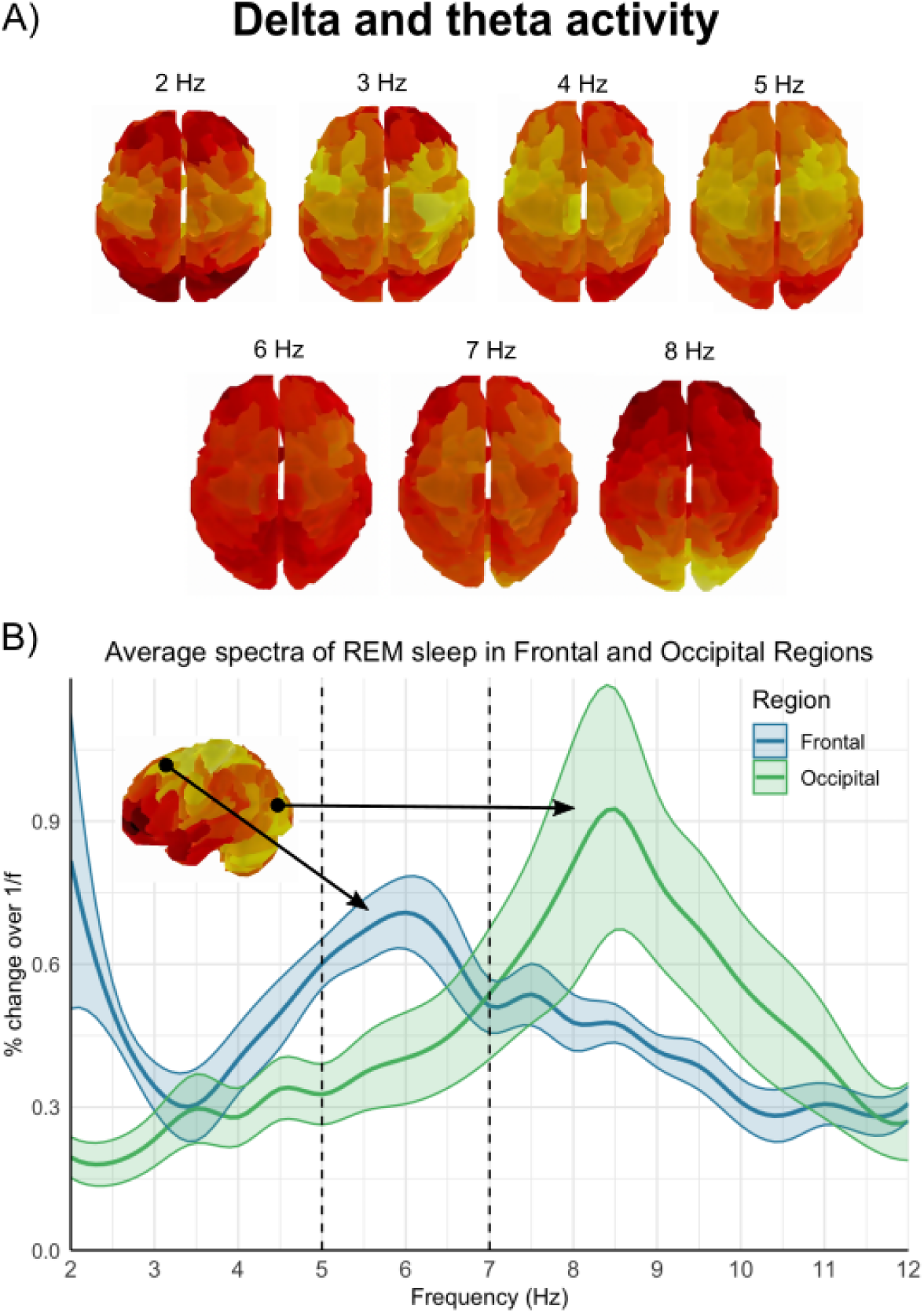
A) Whole-brain topographical maps of the median oscillatory activity across participants for each frequency of the delta and theta band (2-8 Hz; dorsal view) revealed a pronounced oscillatory activity pattern in the occipital regions around 8 Hz. Colour ranges are scaled independently for visualization see, Table 1 in Supplementary Materials for ranges. B) Average spectra plot of activity in REM sleep in a sample frontal and occipital region (regions indicated on a 4-8 Hz topography). The blue and green lines indicate the average spectra of a frontal and occipital region, respectively (shaded error bars = SEM). To reduce the risk of conflating theta and alpha activity, we focused on a narrowed theta range of 5-7 Hz in subsequent analyses (the updated theta frequency range is represented by the dashed lines).

### 4.3 Comparative analysis of theta patterns of REM sleep

#### 4.3.1 Theta oscillations in REM vs. NREM sleep

The total number of retained epochs across subjects, after removing outliers and including only values within 1.5 times the IQR from the first and third quartiles, was 2,027,410 (1,023,705 values for each condition). We identified 261 statistically significant regions (p < 0.05, FDR corrected) with a greater percent change of oscillatory over fractal activity in the (5-7 Hz) theta band during REM sleep compared to NREM sleep (Figure 3A). Topographical mapping revealed prominent theta activity in frontal-central, superior parietal, and temporal regions during REM sleep (for a list of specific regions, see Table 4 in Supplementary Materials). Subcortical regions where theta-band activity was significantly greater than alpha-band activity included the hippocampus, medial septum, amygdala, basal ganglia, thalamus, PPN, and LGN (Figure 3B). No activity was observed in occipital or cerebellar regions, suggesting that the restricted theta-band range (5-7 Hz) was sufficient to remove occipital alpha-like activity in REM sleep. No significant differences were found where NREM sleep exhibited greater theta activity than REM sleep, confirming that in the restricted theta band, as in previous research using the wider frequency range, theta activity is generally stronger in REM than NREM sleep.

**Figure 3:**
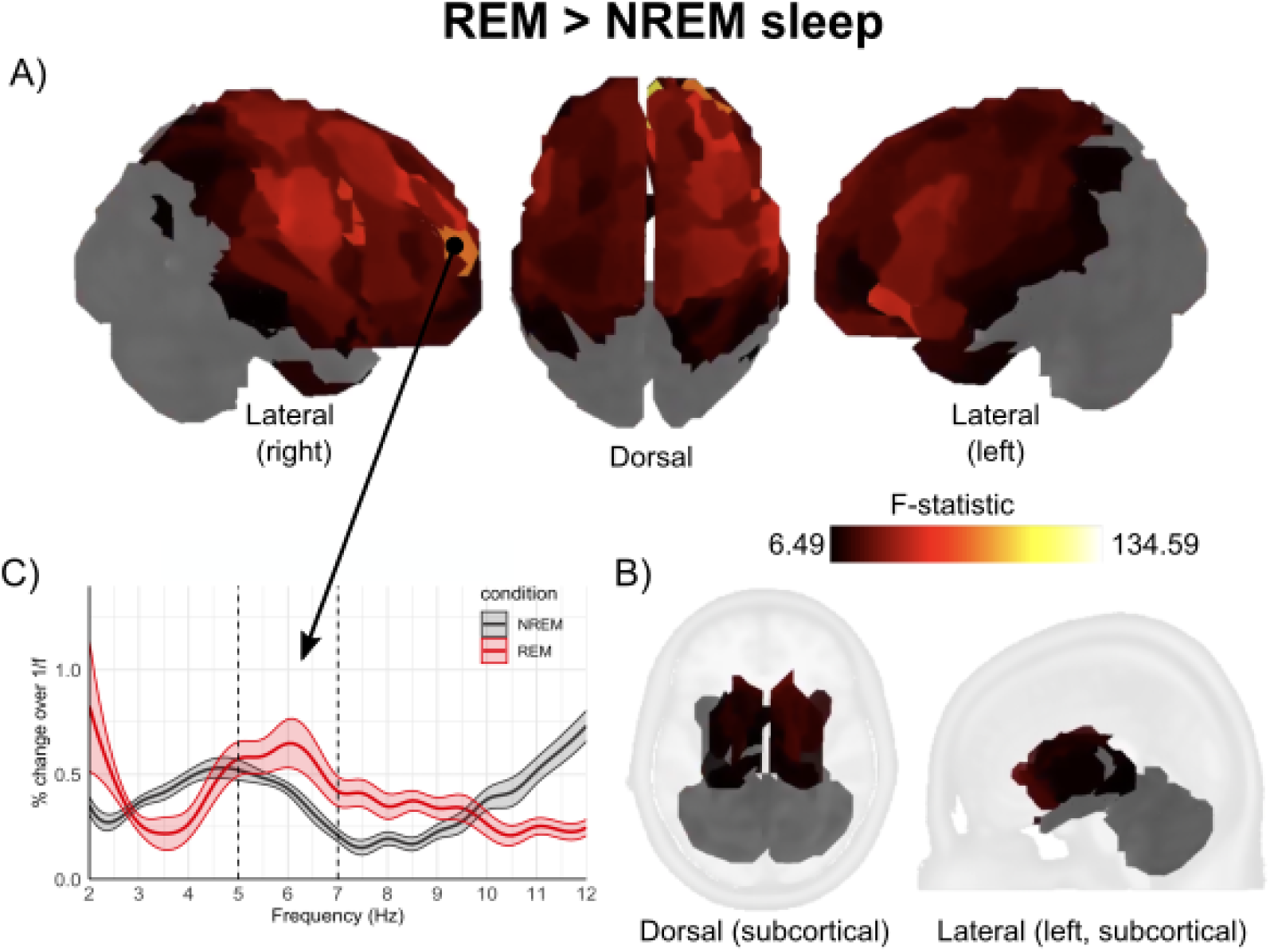
(A) Cortical and (B) subcortical topography of statistically significant regions of interest showing greater theta activity (5-7 Hz) during REM sleep compared to NREM sleep, as determined by LME statistical models (p < 0.05, FDR corrected). No activity in the theta band was observed in the occipital or cerebellar regions, suggesting that the narrow theta-band range was sufficient to remove occipital alpha activity in REM sleep. Once occipital alpha activity was excluded, theta band activity was stronger over frontal and central regions, as well as subcortical regions including the right hippocampus, medial septum, amygdala, basal ganglia (caudate and putamen), thalamus, PPN, and LGN during REM sleep. These findings suggest that activity from these regions in theta could be related to REM’s more cognitive roles. (C) Average spectra plot of activity in REM and NREM sleep stages in an exemplary frontal region. The red and black lines depict the average spectra of REM and NREM sleep stages, respectively (shaded error bars = SEM).

#### 4.3.2 Theta oscillations in phasic vs. tonic REM sleep

The total number of retained epochs across subjects, after removing outliers and including only values within 1.5 times the IQR from the first and third quartiles, was 1,318,128 (659,064 each for each condition). We identified 197 statistically significant regions (p < 0.05, FDR corrected) where the percent change of oscillatory over fractal activity in the theta band was greater during tonic REM sleep compared to phasic REM sleep (Figure 4). Theta activity was stronger in the inferior and lateral posterior regions of the brain, in medial anterior frontal regions (for a detailed list of specific regions, see Table 5 in the Supplementary Materials). In regard to subcortical regions, theta activity was greater in tonic than phasic REM in the medial septum, left hippocampus, left amygdala, basal ganglia, thalamus, PPN, LGN, and the cerebellum. Regions with no significant differences in theta activity between tonic and phasic REM were predominantly located in the superior and medial areas of the brain (note that these regions do show stronger theta activity in the REM vs. NREM comparison, Figure 3). No significant differences were found where phasic REM sleep exhibited greater theta activity than tonic REM sleep.

**Figure 4:**
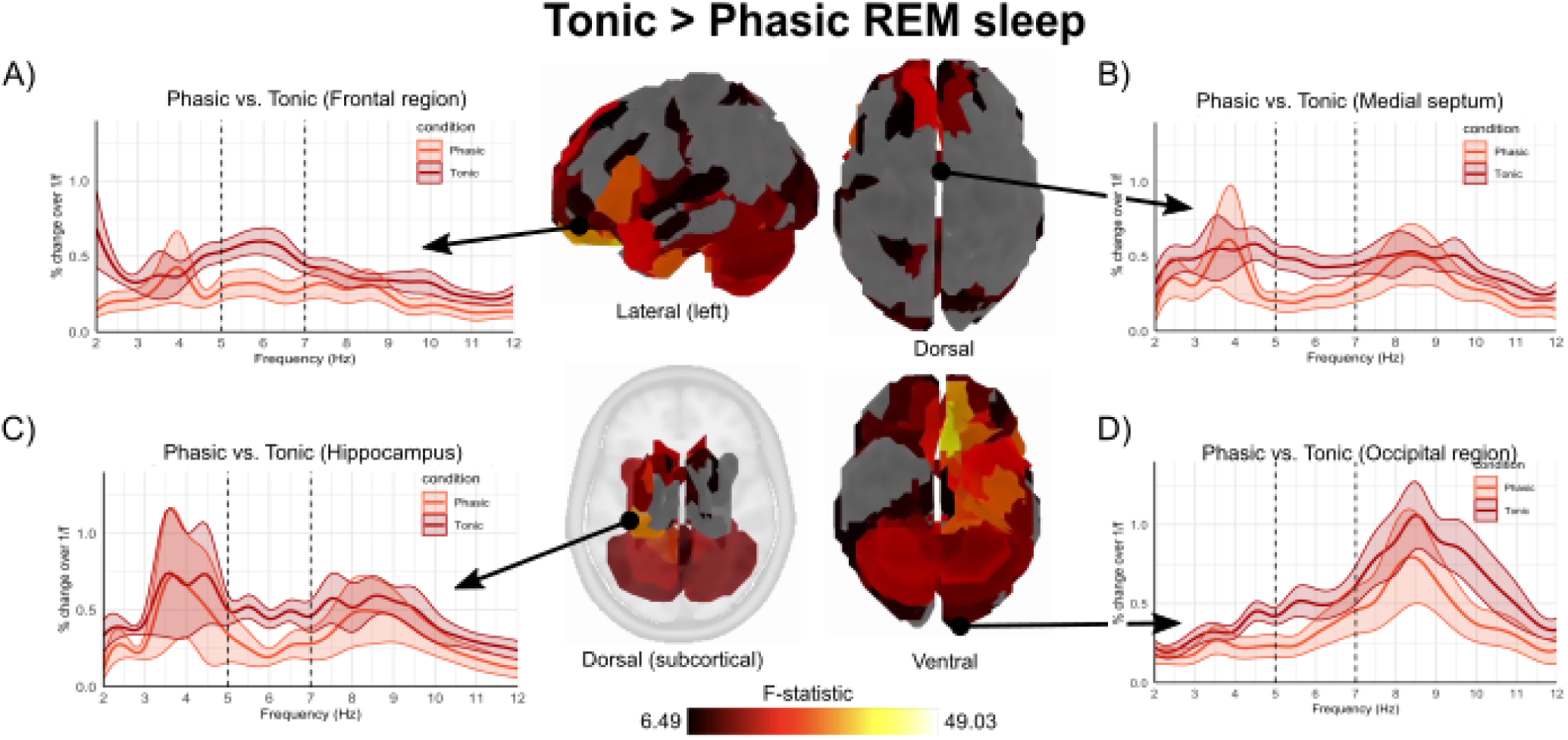
Whole-brain spatial topography revealed statistically significant regions with greater theta activity (5-7 Hz) during tonic REM sleep compared to phasic REM sleep, determined by LME models (p < 0.05, FDR corrected). No significant differences were found where phasic REM sleep exhibited greater activity than tonic REM sleep. Regions showing greater theta activity in tonic REM sleep were located in the inferior and lateral posterior areas of the brain, as well as in a few anterior regions. Subplots A-D) depict average spectra phasic and tonic REM substages in representative frontal, subcortical (i.e., hippocampus and medial septum), and occipital regions (5-7 Hz range is marked with vertical dashed lines). The orange and red lines depict the average spectra of phasic and tonic states, respectively (shaded error bars = SEM).

The overall amplitude of theta activity was consistently higher in tonic states compared to phasic states in widespread brain regions, though notably not in dorsal regions where the theta pattern is overall most prominent in REM sleep (Figure 3). These results suggest that theta activity in tonic sleep is less focal and localized to dorsal regions, rather then being greatly stronger overall. Figure 4)A-D illustrates average spectra from sample regions of interest. Interestingly, although the subcortical regions (C) hippocampus and B) medial septum) do show greater activity in the 5-7 Hz range in tonic REM, the overall spectra appear to show greater distinction between a peak in lower frequencies (3-4 Hz) and in higher frequencies (8-9 Hz) in phasic REM, with the restricted 5-7 Hz theta range being a frequency band of relative inactivity.

### 4.4 REM sleep theta versus working memory theta

To compare theta activity during active wakefulness with theta activity during sleep substages, we first created group (median) topographies for each sleep substage condition. Visual inspection of the topographical map of working memory revealed a distinct pattern of theta activity (5-7 Hz) prominent in the frontal and midline regions of the brain (Figure 5A, left). A similar pattern was observed during phasic REM sleep, but not in other sleep substages, including tonic REM sleep and NREM sleep stages (i.e., N2 K-complex, N2 Spindle, N2 Plain, and SWS). The results of the statistical comparison, using correlation as a measure of similiarity between the mean sleep templates from the sleep dataset with each of the subjects’ topographies in the working memory dateset, indicated that whole-brain theta topographies during the working memory task correlated positively only with phasic REM sleep (V = 155.00, p < 0.001). Instead, they correlated negatively with tonic REM sleep (V = 4.00, p < 0.001) and each NREM sleep stage p < 0.001).

**Figure 5:**
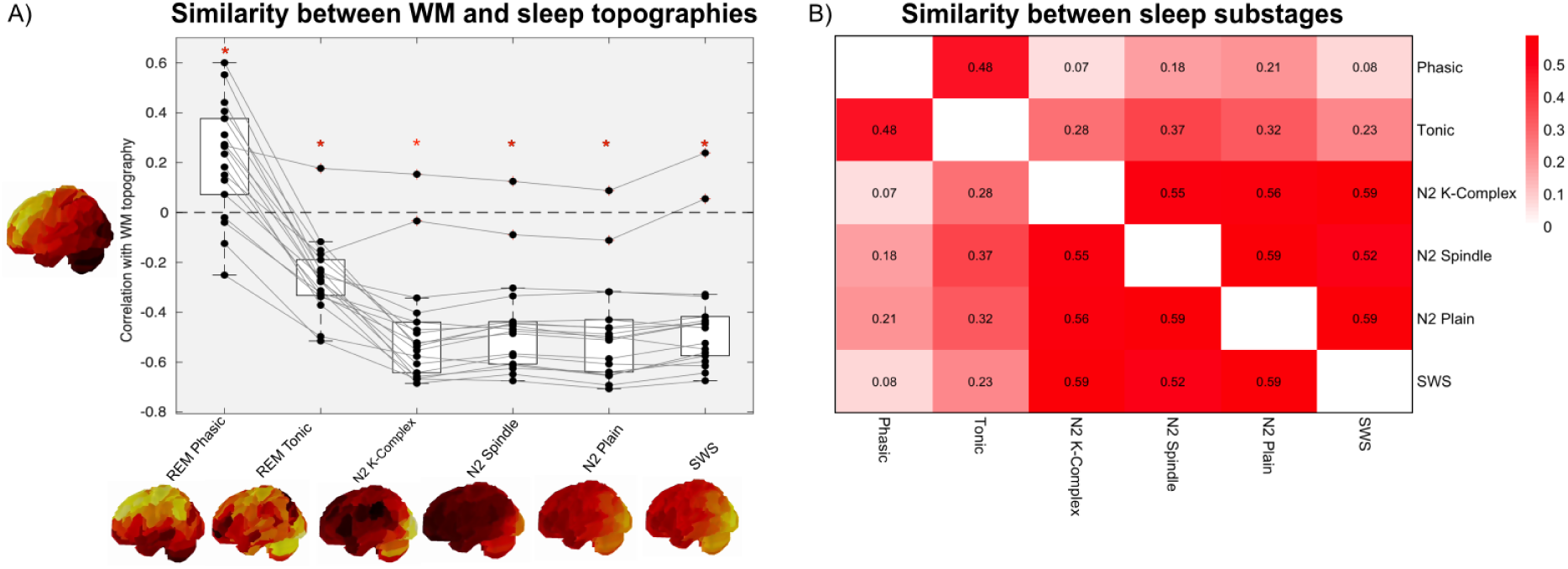
Comparison of theta (5-7 Hz) topographies across states of consciousness. A) Similarity between each subject’s working memory topography (N = 17) and the median topography of each sleep stage across all subjects (N = 10). Asterisks indicate similarity (as measured using correlations) which are significantly different from 0 at the group level (alpha level of 0.01, Wilcoxon signed-rank test). There was a positive correlation between working memory and REM phasic sleep, while a negative correlation was found between working memory and all other sleep stages. Note that brain images are scaled separately to enhance visibility of prominent patterns (see Methods). B) Similarity matrix across sleep substages (N = 10). Mean correlation values are presented for each pair of sleep microstage topographies, averaged across subjects. Tonic and phasic REM are more similar to one another than they are to the NREM substages, which are themselves intercorrelated.

To document relationships between theta topographies across the sleep substages, we present a similarity matrix in which the correlation across sleep substages for each subject averaged across subjects (see Figure 5B). These result further contextualize the apparent dissimilarity in the relationship between phasic vs. tonic results with working memory as presented in Figure 4A. Although phasic and tonic REM sleep substages do differ from one another and phasic REM theta has a relatively stronger similarity to working memory theta, phasic and tonic REM are nevertheless more similar to one another than they are to the NREM sleep substages, which themselves are highly intercorrelated.

## 5 Discussion

In this study, we aimed to provide a spatially-resolved, whole-brain characterisation of REM theta activity in healthy human subjects using MEG. Our findings indicated that (i) theta-band activity has a frontal midline topography that is distinct from those of other frequency bands that are prominent in REM sleep. (ii) Theta activity in the frontal midline regions is best observed by narrowing the theta frequency band to a core range of 5-7 Hz, which allows for better distinction from overlapping alpha-band activity in the occipital and cerebellar regions (Figure 2). (iii) Within this frequency band, theta-band activity during REM sleep as compard to NREM sleep was prominent in frontal and midline cortical structures, but also in subcortical regions (hippocampus, medial septum, amygdala, basal ganglia (caudate and putamen), thalamus, PPN, and LGN; Figure 3). (iv) A closer look into REM sleep substages revealed that theta activity is greater in tonic than phasic REM sleep in inferior and lateral posterior areas but of similar strength over the frontal midline, indicating a less focal frontal midline pattern (rather than an overall increase in theta activity) (Figure 4). (v) Lastly, similarities were observed in whole-brain theta topographies between working memory tasks and phasic REM sleep in frontal midline regions; theta topographies during the working memory task correlated positively with phasic REM sleep, and negatively with all other sleep stages (Figure 5). Tonic and phasic REM theta topographies were however more similar to one another than to those of NREM sleep substages.

### 5.1 Whole-brain oscillatory activity during REM sleep

We aimed to first illustrate the whole-brain topographies of neural oscillations in frequency bands that are prominent in REM (Figure 2A), and then focus on theta-band activity, distinguishing its spatiotemporal patterns from potential overlapping networks in adjacent frequency bands. Activity in the beta frequency range (15-30 Hz) was localized to lateral frontal areas, notably bilaterally dorsolateral and ventrolateral pre-frontal cortex with the strongest signal from the inferior frontal gyrus (which, interestingly, is an area which is the main source of top-down inhibitory signals in task behaviour, e.g., Schaum et al., 2021). Alpha range (8-12 Hz) activity was specific to the occipital regions, and theta range (4-8 Hz) activity was present in frontal midline structures, as well as occipital regions. These topographies are generally consistent with those observed in awake resting state recording using MEG (see Niso et al., 2016)), allowing for some differences possibly due to different frequency ranges (e.g., beta within a more restricted 15-19 Hz range in Niso et al. was more dorsal than lateral), and whether there was task engagement (which also generates a more ventral pattern of activity as seen in Schaum et al., 2021). These similarities might suggest that the same brain regions and circuits which produce oscillatory in particular frequency bands in wakefulness, the functions of which are generally better-characterized, are also present and potentially involved in similar computations in REM sleep. The presence of theta and beta activity in the frontal regions support results from intracranial recordings showing that frontal areas are highly active in REM sleep (Vijayan et al., 2017).

In the present work, we focused on the theta frequency band. The majority of previous EEG studies have compared the amplitude of oscillations averaged within set frequency ranges to quantify the strength of neural activity. Theta and alpha networks during REM sleep likely serve distinct functions, as they do during wakefulness, making it important to be able to separate them (Başar & Güntekin, 2012; Riddle et al., 2020). Magnetoencephalography allows for investigation of topographies by frequency on a fine-grained basis. Our results (Figure 3B) show that the frequency range used in many previous REM sleep studies (i.e., 4-8 Hz) may capture activity from several neural sources, including occipital areas where activity peaks slightly above the theta range (at 8.5 Hz), but has considerable strength at 7-8 Hz; see Figure 2B,C). This frequency overlap could obscure the true spatial patterns and functional roles of theta rhythms during REM sleep, and could make it difficult to target networks with different functions in investigations. A more focused ’core’ range of 5-7 Hz best captured frontal and midline activity during REM sleep, in agreement with previous EEG studies that have observed frontal theta activity (Nishida et al., 2009; Simor et al., 2016; Vijayan et al., 2017). However, even using the focused core theta range, we do observe some occipital activity in some of the other sleep mircostates, notably in tonic REM (Figure 4) and when slow oscillations are present (i.e., N2 K-Complex and SWS NREM substages; Figure 5A). The amplitude and even peak frequency of alpha activity is known to change across states of consciousness including sleep stages and anesthesia (Mierau et al., 2017; Purdon et al., 2015). These findings have methodological implications for research questions that involve comparing neural activity across brain states, as is common in sleep research. Comparing the amplitude of oscillations within frequency bands in the absence of spatial information, even within a restricted range, is likely to capture overlapping networks. In conjunction with previous work suggesting that theta frequency ranges might differ between human and non-human animals (with the human equivalent of theta-band rodent activity being closer to 1-4 Hz; summarised in Buzśaki and Watson, 2012), and that theta frequency might fluctuate over time according to other physiological processes (Bueno-Junior et al., 2023), our results underscore the need for caution in interpreting amplitude in fixed frequency bands rigidly, particularly in the absence of spatial information.

### 5.2 Characterising theta oscillations in REM sleep

Our primary objective was to identify the distinct spatiotemporal patterns of the theta-band during REM sleep, by directly comparing its oscillatory activity to that of other sleep stages, such as NREM. Previous EEG studies have identified oscillatory activity during REM sleep, but these findings are based on data from a limited number of channels, or from EEG, which offers less spatial accuracy (Baillet, 2017; Nishida et al., 2009; Simor et al., 2016). The use of distributed source reconstruction methods in MEG in our study allowed for greater specificity in identifying the spatial distribution of theta activity across the entire brain, including subcortical regions, which are frequently omitted from MEG analyses.

Whole-brain topographies of theta activity during REM sleep revealed a clear pattern of oscillations in the frontal lobe. Our results align with previous EEG and intracranial studies that have observed theta activity in frontal regions such as the dlPFC and ACC during REM sleep (Nishida et al., 2009; Simor et al., 2016; Vijayan et al., 2017). Our results extend these findings by identifying theta rhythms across a wider range of frontal regions, including inferior, medial, middle, and superior frontal regions. This widespread distribution of theta activity may suggest a more integrated role for the frontal lobe in the cognitive processes associated with REM sleep, such as memory consolidation.

Previous PET studies have reported hypoactivation of the dlPFC during REM sleep compared to NREM sleep (Braun et al., 1997; Maquet et al., 2005). However, neural synchronization may not have a direct, positive relationship with energy consumption (Scheeringa et al., 2011), meaning that decreased activation in PET or fMRI signals is not necessarily at odds with the observation of increased theta-band activity. Results from an intracranial EEG study have in fact revealed theta-band activity in the dlPFC during REM sleep (Vijayan et al., 2017). The results of our study corroborate these findings, revealing that theta activity in the dlPFC is greater during REM sleep, as compared to NREM sleep. Our findings, along with those of Vijayan et al., 2017, suggest that the dlPFC plays a more significant role in REM sleep than previously understood. During wakefulness, the dlPFC is involved in the maintenance and manipulation of information during working memory tasks (Barbey et al., 2013; Blumenfeld & Ranganath, 2006). For example, fMRI studies showed dlPFC activation during working memory tasks that require chunking information into smaller units and rearranging the order of items (Blumenfeld & Ranganath, 2006; Bor et al., 2004). During REM sleep, the dlPFC may play related roles, for example helping to reorganize neural representations of experiences to facilitate their integration with existing knowledge. Although it is intriguing to consider that the dlPFC may be involved in certain cognitive processes related to memory or dreaming, these physiological results do not directly inform us as to their functional roles in REM sleep, for which further research will be required.

Our study corroborates previous PET studies identifying the neural substrates active during REM sleep, while extending these findings by using MEG to reveal that these regions are specifically oscillating within the theta band. Specifically, our results replicate previous PET studies comparing REM to NREM sleep by identifying theta activity in regions such as the anterior cingulate cortex, frontal medial cortex, fusiform gyrus, insula, parahippocampal gyrus, precuneus, supplementary motor area, temporal gyrus, and temporal pole (Braun et al., 1997; Maquet et al., 2005). While the precise functional role of these theta oscillations remains to be understood, from a cognitive perspective, this widespread activity may support the integration of certain cognitive processes, potentially facilitating memory consolidation and the integration of new learning with prior knowledge, and/or other higher-order brain functions during REM sleep.

Investigating deep subcortical sources using MEG is a relatively new method. Initially, it was believed that MEG could only effectively localise cortical sources because deeper sources produce signals that are more attenuated by their central location relative to the sensor helmet and distance from source to sensors (Hämäläinen et al., 1993). However, recent advances in methods have improved sensitivity to deep brain sources, including those in the cerebellum, thalamus, and brainstem (Andersen et al., 2020; Coffey et al., 2016, 2021). Our results demonstrate that it is possible to record theta activity during REM sleep in subcortical structures, including the amygdala, basal ganglia, hippocampus, medial septum, PPN, LGN, and thalamus (however noting that the ability of these methods to resolve activity from proximal deep sources is still under study). Most of the regions where we observed theta activity are known to be crucial for REM sleep. For example, the hippocampus and medial septum are essential for memory processing and the synchronisation of theta rhythms (Boyce et al., 2016). The hippocampus and amygdala play important roles in the expression of drive and affect, as well as in the control of autonomic function (Braun et al., 1997). The PPN and thalamus are involved in REM initiation and maintenance, contributing to the generation and modulation of REM sleep-associated neural activity (Urbano et al., 2014). The LGN is associated with processing visual information during REM sleep (particularly in relation to ponto-geniculo-occipital (PGO) waves), while the basal ganglia are involved in motor control and the regulation of sleep-wake transitions (Braun et al., 1997; Steriade et al., 1989). The results presented here highlight MEG’s potential as a tool for exploring spatiotemporal patterns across the entire brain in healthy humans and further investigating the functional contributions of subcortical brain regions across brain states.

Examining REM sleep substages more closely, we found that theta oscillations in tonic REM appear more prominently in inferior and lateral posterior areas than they do in phasic REM. This result is best interpreted in conjunction with the REM vs. NREM result (Figure 3) and with reference to the similarity analyses displayed in Figure 5: REM in general as compared with NREM is characterised by theta-band activity over frontal and dorsal cortical structures, and tonic and phasic REM sleep topographies are quite highly correlated (i.e., r = 0.48; Figure 5B), yet oscillatory activity is less regionally-concentrated in tonic vs. phasic substages, with theta in tonic REM observed in wider selection of ventral regions. Extracting spectra from representative regions during each microstate offers further insight: in both a frontal cortical region (Figure 4) and an occipital region, tonic REM has slightly higher oscillatory activity in the 5-7 Hz range than does phasic REM. In some subcortical regions (medial septum, hippocampus), theta activity appears to have two peaks; one at 2-4 Hz and the other at 8-9 Hz; this pattern is more distinct in phasic REM (in accordance with previous work, Simor et al., 2020), whereas tonic REM exhibited somewhat elevated activity across the intervening range, which includes our 5-7 Hz focus.

These results support the idea that tonic and phasic periods in REM state are distinct states (Simor et al., 2020). Although in previous work EEG strength averaged at the scalp activity in the 5-7 Hz was found to be greater in phasic than tonic REM (Simor et al., 2020), these results are likely accounted for by differences in methodological approaches; in Simor et al. the metric was global average of EEG signals in sensor space, without adjusting for microstate differences in background aperiodic signal. The present results use source-localization in MEG signals, and separate oscillatory activity from fractal activity (which can change across brain state including phasic and tonic REM; Lendner et al., 2020; Rosenblum et al., 2024; Wen and Liu, 2016). The present results demonstrate that MEG can be used to observe fine-grained patterns in frequency of subcortical brain regions, and provide suggestions for where to look for further differences between substages. For example, although the phasic vs. tonic REM statistical comparison did not reveal any brain regions in which phasic REM had greater activity in the 5-7 Hz range, extracted spectra across a broader range of frequencies from cortical and subcortical sources (Figure 4A-D) hint that a special feature of phasic REM might actually be 2-4 Hz activity rather than theta, and that it might be generated in subcortical regions including hippocampus and medial septum. As these regions are causally implicated in memory consolidation in animal models (Boyce et al., 2016), and the equivalent neural functions may take place in a lower frequency range in humans as compared with rodents (Bódizs et al., 2001; Buzśaki & Watson, 2012), it is possible that this lower frequency band (potentially coupled to gamma-band activity) is involved in coordinating transfer of spatial and temporally-organised information with the hippocampus (Nuñez & Buño, 2021), whereas cortical theta over a broader frequency range is engaged in other processes such as integrating the information with previous knowledge. Visually comparing the delta and theta topographies presented in Figure 1 also suggests that this idea merits further investigation; in the delta band there is more extensive subcortical activation, including hippocampus.

### 5.3 Similarities and dissimiliarties between REM theta and working memory theta

We observed a high similarity in theta topography between phasic REM sleep and waking working memory; both showed a pattern of stronger activity in frontal midline areas (Figure 5). This similarity was observed only for phasic REM sleep. The working memory pattern correlated negatively with all other sleep substages. The exclusive similarity observed between theta activity during phasic REM sleep and working memory tasks in wakefulness suggests that a frontal theta network may serve similar functions in both states, and that network activity might be suppressed in NREM sleep, in which only the occipital alpha-like pattern was observed.

Theta activity associated with working memory processes has previously been observed in frontal midline regions (Hsieh & Ranganath, 2014). In sleep, external stimuli and environmental alertness is relatively preserved in tonic REM sleep, and is decreased in phasic REM sleep (Simor et al., 2020). This distinction suggests that phasic REM sleep serves as an internally-focused processing state, potentially facilitating complex cognitive processes. Given that phasic REM sleep appears to support internal cognitive processes and that theta activity has a focused pattern over frontal midline structures during this microstate, it could be an important period for activities related to cognitive processes including memory consolidation.

In memory models of sleep, the Sequential Hypothesis of Memory Consolidation suggests that NREM sleep strengthens individual memory traces, while REM sleep integrates these traces into existing knowledge networks (Giuditta, 2014). The dlPFC plays a crucial role in maintaining and manipulating information during working memory tasks (Barbey et al., 2013; Blumenfeld & Ranganath, 2006). The presence of theta oscillations in the dlPFC during phasic REM sleep suggests that similar mechanisms might be at play. For example, theta activity in frontal regions may be critical for reorganising neural representations of experiences and integrating new information with existing memories. The involvement of the dlPFC in these processes during REM sleep could aid in refining and stabilising memory traces. During REM sleep, the activation of neural networks similar to those involved in working memory may promote the reorganisation of neural representations. This reorganisation may support the integration of new information with pre-existing memories, thereby enhancing the overall consolidation process.

Interestingly, tonic REM topographies were negatively correlated with working memory theta (although noting that the inter-correlation between phasic and tonic REM stages was somewhat positively correlated, Figure 5B). This pattern of results suggests that whilst there are general similarities in theta activity between these substages, there are also differences in how they relate to waking activity, and therefore potentially in which roles they serve in sleep-dependent information processing. This line of questioning may eventually address why REM sleep exists; if some aspects of REM physiology and function are not unique to the sleep state, why not stay awake? Some hints may come from investigations of cross-frequency coupling. In a mouse model, Scheffzük et al., 2011 showed that interactions between theta and fast gamma activity (120-160 Hz) increased 9-fold during REM as compared to active wakefulness (noting that phasic and tonic sleep substages were not distinguished in the mouse model). While we do not have direct insight into differential functional roles through the present physiological analysis, the results encourage further work differentiating tonic and phasic REM (Simor et al., 2020), through fine-grained analysis of oscillatory activity.

### 5.4 Limitations and future work

Our sample size for the sleep analysis is relatively small (N = 10 healthy young adults). To address this limitation, we employed linear mixed-effects models (LMEs) to account for the nested structure of our data, which includes thousands of epochs per subject. This statistical approach is particularly advantageous for our study design, as it allowed us to capitalise on the extensive data from each subject, thereby enhancing the robustness and validity of the results (Schielzeth et al., 2020). Furthermore, our study focuses specifically on the electrophysiological features of sleep, which generally exhibits less variability compared to behavioural studies. While the study’s physiological focus is a strength as regards obtaining high SNR within a small sample, the study design is of a passive, observational nature; we have not manipulated REM theta, and so our interpretation as regard theta function similar remain correlational and speculative. We nonetheless believe that characterising REM theta in healthy humans is an important and necessary precursor to these more mechanistic investigations.

The analyses conducted in the present work used measures of oscillatory activity to look at whole-brain spatial topographies. This approach has the advantage of being robust and relatively straightforward (i.e., few parameters to choose and producing results that are consistent with previous work), but does not capture communication between brain regions (i.e., functional connectivity), and does not take advantage of that potentially valuable information to disentangle networks that overlap spatially or temporally. In part this design choice was made as methods for characterizing functional connectivity particularly in three dimensional deep source models require further methodological development and validation prior to being applied. In the future, methods that can isolate and examine brain networks may obviate the need for defining frequency ranges and allow for greater insight into network networks and circuits sub-serving memory functions in REM sleep (e.g., Pelzer et al., 2024; Yu et al., 2022).

## 6 Conclusion

Using MEG, we were able to capture the frequency-specific activity of brain regions during REM sleep, identifying the spatiotemporal patterns of theta oscillations, including in deep subcortical sources. These findings can help guide future studies in several ways. Future research can use closed-loop auditory stimulation (CLAS) and other brain stimulation techniques to offer causal insights into the role of theta oscillations in REM sleep. For instance, subsequent experimental studies could target prefrontal theta activity using CLAS to modulate and enhance memory consolidation processes. By applying stimulation during REM sleep when theta activity is naturally occurring, researchers could investigate how altering theta rhythms impacts memory consolidation (Harrington et al., 2021). Subsequent investigations using MEG should investigate other frequency bands of interest in REM sleep, including delta, beta and gamma oscillations, as well as interactions between freqency bands. Theta-gamma coupling may be particularly interesting, as it has been observed during both REM sleep and working memory tasks in the hippocampus and neocortex (Bandarabadi et al., 2019; Scheffzük et al., 2011; Tamura et al., 2017), and may be most pronounced in REM sleep (Scheffzük et al., 2011). A better understanding of this coupling might reveal REM’s unique roles in cognition. Lastly, there is limited research on the cerebellum’s role in REM sleep (see Canto et al., 2017 for a review). We observed theta activity within the cerebellum during REM sleep, aligning with a previous study that reported cerebellar activation (Braun et al., 1997). The cerebellum’s role in sleep has recently started to gain interest after the discovery that it generates sleep spindles, which are hallmarks of memory consolidation in NREM sleep (Xu et al., 2021). Our whole-brain topographies hint at divergent patterns of activity in the cerebellum across frequency bands (Figure 1) and sleep states (Figure 5) further supporting the notion that MEG can be used to study cerebellar activity (Andersen et al., 2020), and open new possibilities to explore the cerebellum’s role in sleep and memory (Benarroch, 2023; Jackson & Xu, 2023). There is much to be done to precisely define the role of REM sleep and its underlying neural mechanisms in sleep-dependent memory consolidation. This work contributes a whole-brain characterisation of the spatiotemporal patterns of theta rhythms in healthy humans to these efforts.

## Data and Code Availability

Extracted values used for statistical analyses are available on the Open Science Framework website (https://osf.io/tk4ry/?view only=b7b8790bcfef4b3283648e111636011b). The raw MEG sleep recordings and working memory task recordings were collected under the supervision of SG at the University of Tübingen and PA at Laval University, respectively. These data are not publicly available due to local ethics restrictions, but may be made accessible pending approval from the principal investigators and local ethics committee.

## Author Contributions

**Vasiliki Provias:** Conceptualisation; Methodology; Formal analysis; Software; Visualisation; Writing—original draft; Writing—review and editing. **Monika Schönauer:** Conceptualisation; Writing—review and editing. **Steffan Gais:** Data collection; Conceptualisation; Resources; Funding acquisition; Writing—review and editing. **Jan Born:** Conceptualisation; Resources; Funding acquisition; Supervision; Writing—review and editing. **Til O. Bergmann:** Conceptualisation; Methodology; Supervision; Writing—review and editing. **Emily B.J. Coffey**: Conceptualisation; Resources; Methodology; Formal analysis; Data curation; Writing—original draft; Writing—review and editing; Supervision; Project administration; Funding acquisition.

## Acknowledgements

We would like to thank all the trainees who collected the data that were used in these analyses. The sleep dataset was provided by Steffen Gais’ lab, acquired by Lea Himmer, Zóe Bürger, Leonie Fresz, Janina Maschke, and Lore Wagner, and the working memory dataset was provided by Philippe Albouy’s lab. We also thank Alberto Romero Ara and Hugo Jourde for their guidance in applying linear mixed-effects models in our analyses, and Hugo Jourde, Chris Steele and Thahn Dang-Vu for helpful comments on an earlier draft. VP was supported by a scholarship from the Quebec Bio-Imaging Network. TOB received funding from the German Research Foundation (DFG) via FOR 5434 ”Information Abstraction During Sleep” (Grant no. 468645090). EBJC was funded by a Banting Postdoctoral Fellowship during early stages of this work and later by a Discovery Grant from the Natural Sciences and Engineering Research Council of Canada (NSERC).

