## Supplementary Material for "Spatiotemporal patterns of theta-band activity during rapid-eye movement sleep: a magnetoencephalography analysis"

### Supplementary Materials

**Figure 1.** Characterising REM theta by comparing to alpha (8-12 Hz)

To robustly evaluate differences in theta versus alpha band activity during REM sleep, we conducted LME analysis at the epoch-trial level, using subjects as a random effect. The total number of retained epochs across subjects, after removing outliers and including only values within 1.5 times the IQR from the first and third quartiles, was 2,642,776 (1,321,388 each for theta and alpha). Our analysis identified 276 regions with statistically significant differences in which theta activity was higher than alpha during REM sleep ($p < 0.05$, FDR corrected). Increased activity in the theta band was observed primarily in frontal, central, temporal, and superior parietal regions. Subcortical regions where theta-band activity was significantly greater than alpha-band activity included the hippocampus, medial septum, amygdala, basal ganglia, thalamus, PPN, and LGN. Subsequently, our analysis identified 106 regions with statistically significant differences in which alpha activity was higher than theta during REM sleep ($p < 0.05$, FDR corrected). Alpha activity is predominantly located in occipital regions, including the striate and extrastriate cortices. In addition, the cerebellum, including the vermis, exhibited significantly higher alpha activity compared to theta activity.

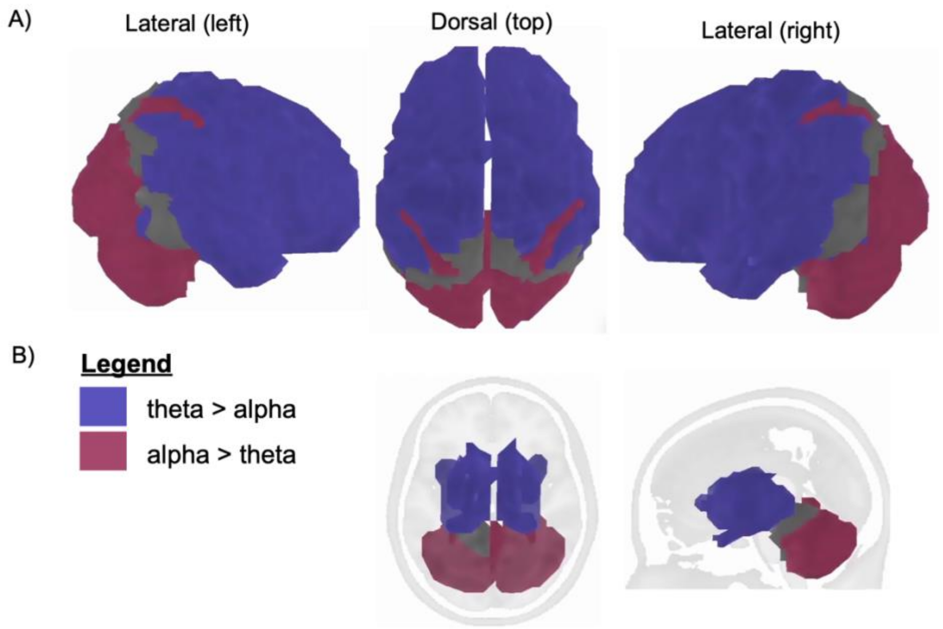


Figure 1. Binarized whole-brain spatial topography of statistically significant F-statistic values (derived from LMEs). Purple regions indicate a higher percentage change in oscillatory over fractal activity within the theta band (5-Hz) compared to the alpha band (8-12 Hz) during REM sleep. Oscillatory activity in the theta band during REM sleep is primarily localised to the frontal, central, temporal, and parietal regions. The hippocampus, medial septum, amygdala,

basal ganglia (caudate and putamen), thalamus, PPN, and LGN were identified as subcortical regions where theta-band activity was significantly greater than alpha-band activity. Conversely, pink regions denote a greater percentage change in the alpha band relative to the theta band. Alpha activity is predominantly located in posterior regions of the brain, such as the striate (primary visual cortex), extrastriate cortices, and the cerebellum (including vermis).}

**Supplementary Table 1.** Maximum and minimum median values during REM sleep within each sliced frequency bin of the 4-8 Hz range (from Figure 1).

| **Sliced frequency bins** | **Minimum value** | **Maximum value** |
|---:|---|---|
| 2 Hz | 0.22 | 0.40 |
| 3 Hz | 0.30 | 0.58 |
| 4 Hz | 0.27 | 0.55 |
| 5 Hz | 0.22 | 0.64 |
| 6 Hz | 0.15 | 0.60 |
| 7 Hz | 0.09 | 0.62 |
| 8 Hz | 0.39 | 0.73 |

**Supplementary Table 2.** Maximum and minimum median values in all sleep stages at 5-7 Hz and during the working memory task (from Figure 4).

| **Theta 5-7 Hz** | **Minimum value** | **Maximum value** |
|---:|---|---|
| Phasic REM sleep | 0.32 | 0.62 |
| Tonic REM sleep | 0.38 | 0.70 |
| N2 K-complex | 0.28 | 0.53 |
| N2 Spindle | 0.26 | 0.50 |
| N2 Plain | 0.23 | 0.58 |
| SWS Deep | 0.27 | 0.59 |
| Working Memory | 0.36 | 0.65 |

**Supplementary Table 3.** Statistically significant regions of interest showing a higher percent change in oscillatory over fractal activity within the theta band (5-7 Hz) compared to the alpha band (8-12 Hz). The table also indicates regions with a greater percent change in the alpha band relative to the theta band. P-values have been corrected for FDR ($p < 0.05$).

| **ROI** | **F-statistic** | **FDR p-value** |
|---:|---|---|
| **G_Angular-1 L** | 9.25 | 0.022 |
| **G_Angular-1 R** | 10.75 | 0.016 |
| **G_Angular-3 R** | 8.27 | 0.029 |
| **G_Cingulum_Ant-1 L** | 37.01 | 0.001 |
| **G_Cingulum_Ant-1 R** | 73.94 | 0.001 |
| **G_Cingulum_Ant-2 L** | 27.68 | 0.002 |
| **G_Cingulum_Ant-2 R** | 38.70 | 0.001 |

| | | |
|---|---|---|
| **G_Cingulum_Mid-1 L** | 19.00 | 0.004 |
| **G_Cingulum_Mid-1 R** | 18.69 | 0.004 |
| **G_Cingulum_Mid-2 L** | 21.68 | 0.003 |
| **G_Cingulum_Mid-2 R** | 18.08 | 0.004 |
| **G_Cingulum_Mid-3 L** | 16.28 | 0.005 |
| **G_Cingulum_Mid-3 R** | 40.13 | 0.001 |
| **G_Cingulum_Post-1 L** | 14.60 | 0.007 |
| **G_Cingulum_Post-1 R** | 22.56 | 0.002 |
| **G_Cingulum_Post-2 L** | 9.49 | 0.021 |
| **G_Cingulum_Post-2 R** | 10.17 | 0.018 |
| **G_Frontal_Inf_Orb-1 L** | 77.54 | 0.001 |
| **G_Frontal_Inf_Orb-1 R** | 25.52 | 0.002 |
| **G_Frontal_Inf_Orb-2 L** | 21.06 | 0.003 |
| **G_Frontal_Inf_Orb-2 R** | 14.20 | 0.007 |
| **G_Frontal_Inf_Tri-1 L** | 43.06 | 0.001 |
| **G_Frontal_Inf_Tri-1 R** | 35.09 | 0.001 |
| **G_Frontal_Med_Orb-1 L** | 25.80 | 0.002 |
| **G_Frontal_Med_Orb-1 R** | 23.05 | 0.002 |
| **G_Frontal_Med_Orb-2 L** | 27.39 | 0.002 |
| **G_Frontal_Med_Orb-2 R** | 29.60 | 0.002 |
| **G_Frontal_Mid_Orb-1 L** | 12.45 | 0.011 |
| **G_Frontal_Mid_Orb-1 R** | 24.97 | 0.002 |
| **G_Frontal_Mid_Orb-2 L** | 10.51 | 0.016 |
| **G_Frontal_Mid_Orb-2 R** | 30.52 | 0.001 |
| **G_Frontal_Mid-1 L** | 28.16 | 0.002 |
| **G_Frontal_Mid-1 R** | 29.75 | 0.001 |
| **G_Frontal_Mid-2 L** | 29.00 | 0.001 |
| **G_Frontal_Mid-2 R** | 41.89 | 0.001 |
| **G_Frontal_Mid-3 L** | 29.19 | 0.001 |
| **G_Frontal_Mid-3 R** | 67.76 | 0.001 |
| **G_Frontal_Mid-4 L** | 34.37 | 0.001 |
| **G_Frontal_Mid-4 R** | 49.43 | 0.001 |
| **G_Frontal_Mid-5 L** | 32.41 | 0.001 |
| **G_Frontal_Mid-5 R** | 46.88 | 0.001 |
| **G_Frontal_Sup_Medial-1 L** | 37.51 | 0.001 |
| **G_Frontal_Sup_Medial-1 R** | 134.59 | 0.000 |
| **G_Frontal_Sup_Medial-2 L** | 36.32 | 0.001 |
| **G_Frontal_Sup_Medial-2 R** | 43.43 | 0.001 |
| **G_Frontal_Sup_Medial-3 L** | 30.03 | 0.001 |
| **G_Frontal_Sup_Medial-3 R** | 29.65 | 0.001 |
| **G_Frontal_Sup_Orb-1 L** | 34.56 | 0.001 |

| | | |
|---|---|---|
| **G_Frontal_Sup_Orb-1 R** | 21.56 | 0.003 |
| **G_Frontal_Sup-1 L** | 33.46 | 0.001 |
| **G_Frontal_Sup-1 R** | 44.36 | 0.001 |
| **G_Frontal_Sup-2 L** | 36.85 | 0.001 |
| **G_Frontal_Sup-2 R** | 36.68 | 0.001 |
| **G_Frontal_Sup-3 L** | 21.36 | 0.003 |
| **G_Frontal_Sup-3 R** | 36.47 | 0.001 |
| **G_Fusiform-1 R** | 7.56 | 0.034 |
| **G_Hippocampus-1 R** | 19.35 | 0.003 |
| **G_Hippocampus-2 L** | 6.64 | 0.044 |
| **G_Hippocampus-2 R** | 11.34 | 0.014 |
| **G_Insula-anterior-1 L** | 11.01 | 0.015 |
| **G_Insula-anterior-1 R** | 7.96 | 0.031 |
| **G_Insula-anterior-2 L** | 28.78 | 0.002 |
| **G_Insula-anterior-2 R** | 18.09 | 0.004 |
| **G_Insula-anterior-3 L** | 29.43 | 0.001 |
| **G_Insula-anterior-3 R** | 42.84 | 0.001 |
| **G_Insula-anterior-4 L** | 30.69 | 0.001 |
| **G_Insula-anterior-4 R** | 21.52 | 0.003 |
| **G_Insula-anterior-5 L** | 21.96 | 0.003 |
| **G_Insula-anterior-5 R** | 38.67 | 0.001 |
| **G_Insula-posterior-1 L** | 16.11 | 0.005 |
| **G_Insula-posterior-1 R** | 25.72 | 0.002 |
| **G_Lingual-1 L** | 6.50 | 0.046 |
| **G_Paracentral_Lobule-1 L** | 28.09 | 0.002 |
| **G_Paracentral_Lobule-1 R** | 31.99 | 0.001 |
| **G_Paracentral_Lobule-2 L** | 34.31 | 0.001 |
| **G_Paracentral_Lobule-2 R** | 38.76 | 0.001 |
| **G_Paracentral_Lobule-3 L** | 30.39 | 0.001 |
| **G_Paracentral_Lobule-3 R** | 31.83 | 0.001 |
| **G_Paracentral_Lobule-4 L** | 29.19 | 0.001 |
| **G_Paracentral_Lobule-4 R** | 33.95 | 0.001 |
| **G_ParaHippocampal-1 L** | 7.06 | 0.039 |
| **G_ParaHippocampal-1 R** | 13.74 | 0.009 |
| **G_ParaHippocampal-2 R** | 7.44 | 0.035 |
| **G_ParaHippocampal-3 L** | 8.13 | 0.030 |
| **G_ParaHippocampal-3 R** | 9.34 | 0.021 |
| **G_ParaHippocampal-4 R** | 11.34 | 0.014 |
| **G_Parietal_Inf-1 L** | 10.67 | 0.016 |
| **G_Parietal_Inf-1 R** | 15.40 | 0.006 |
| **G_Parietal_Sup-1 L** | 16.91 | 0.005 |

| | | |
|---|---|---|
| **G_Parietal_Sup-1 R** | 16.35 | 0.005 |
| **G_Parietal_Sup-2 L** | 18.85 | 0.004 |
| **G_Parietal_Sup-2 R** | 17.20 | 0.005 |
| **G_Parietal_Sup-3 L** | 21.49 | 0.003 |
| **G_Parietal_Sup-3 R** | 10.74 | 0.016 |
| **G_Parietal_Sup-4 L** | 10.71 | 0.016 |
| **G_Parietal_Sup-4 R** | 16.05 | 0.006 |
| **G_Parietal_Sup-5 R** | 9.12 | 0.023 |
| **G_Precuneus-4 R** | 8.21 | 0.030 |
| **G_Precuneus-5 R** | 13.00 | 0.011 |
| **G_Rolandic_Oper-1 L** | 32.70 | 0.001 |
| **G_Rolandic_Oper-1 R** | 26.88 | 0.002 |
| **G_Rolandic_Oper-2 L** | 26.95 | 0.002 |
| **G_Rolandic_Oper-2 R** | 47.62 | 0.001 |
| **G_subcallosal-1 L** | 18.16 | 0.004 |
| **G_subcallosal-1 R** | 22.31 | 0.002 |
| **G_Supp_Motor_Area-1 L** | 26.58 | 0.002 |
| **G_Supp_Motor_Area-1 R** | 39.41 | 0.001 |
| **G_Supp_Motor_Area-2 L** | 28.33 | 0.002 |
| **G_Supp_Motor_Area-2 R** | 36.74 | 0.001 |
| **G_Supp_Motor_Area-3 L** | 34.10 | 0.001 |
| **G_Supp_Motor_Area-3 R** | 30.83 | 0.001 |
| **G_Supramarginal-1 L** | 21.38 | 0.003 |
| **G_Supramarginal-1 R** | 37.36 | 0.001 |
| **G_SupraMarginal-2 L** | 22.76 | 0.002 |
| **G_SupraMarginal-2 R** | 34.57 | 0.001 |
| **G_Supramarginal-3 L** | 34.78 | 0.001 |
| **G_Supramarginal-3 R** | 56.98 | 0.001 |
| **G_Supramarginal-4 L** | 27.11 | 0.002 |
| **G_Supramarginal-4 R** | 21.73 | 0.003 |
| **G_SupraMarginal-5 L** | 19.14 | 0.004 |
| **G_SupraMarginal-5 R** | 18.57 | 0.004 |
| **G_SupraMarginal-6 L** | 16.32 | 0.005 |
| **G_SupraMarginal-6 R** | 13.79 | 0.008 |
| **G_SupraMarginal-7 L** | 9.87 | 0.019 |
| **G_SupraMarginal-7 R** | 15.31 | 0.006 |
| **G_Temporal_Inf-1 L** | 10.53 | 0.017 |
| **G_Temporal_Inf-1 R** | 16.88 | 0.006 |
| **G_Temporal_Mid-1 L** | 10.04 | 0.019 |
| **G_Temporal_Mid-1 R** | 17.54 | 0.005 |
| **G_Temporal_Mid-2 R** | 6.83 | 0.042 |

| | | |
|---|---|---|
| **G_Temporal_Mid-3 L** | 10.14 | 0.018 |
| **G_Temporal_Mid-3 R** | 13.27 | 0.010 |
| **G_Temporal_Pole_Mid-1 L** | 17.57 | 0.004 |
| **G_Temporal_Pole_Mid-2 L** | 11.98 | 0.012 |
| **G_Temporal_Pole_Mid-3 L** | 7.76 | 0.032 |
| **G_Temporal_Pole_Mid-3 R** | 17.58 | 0.004 |
| **G_Temporal_Pole_Sup-1 L** | 31.59 | 0.001 |
| **G_Temporal_Pole_Sup-1 R** | 7.74 | 0.032 |
| **G_Temporal_Pole_Sup-2 L** | 22.74 | 0.002 |
| **G_Temporal_Pole_Sup-2 R** | 9.56 | 0.020 |
| **G_Temporal_Sup-1 L** | 26.43 | 0.002 |
| **G_Temporal_Sup-1 R** | 43.44 | 0.001 |
| **G_Temporal_Sup-2 L** | 13.73 | 0.009 |
| **G_Temporal_Sup-2 R** | 25.68 | 0.002 |
| **G_Temporal_Sup-3 L** | 17.91 | 0.004 |
| **G_Temporal_Sup-3 R** | 24.95 | 0.002 |
| **G_Temporal_Sup-4 L** | 16.55 | 0.005 |
| **G_Temporal_Sup-4 R** | 33.88 | 0.001 |
| **LGNleft** | 7.53 | 0.034 |
| **LGNright** | 13.37 | 0.009 |
| **medsep** | 9.82 | 0.019 |
| **N_Amygdala-1 L** | 7.25 | 0.037 |
| **N_Amygdala-1 R** | 9.48 | 0.020 |
| **N_Caudate-1 L** | 19.85 | 0.003 |
| **N_Caudate-1 R** | 24.16 | 0.002 |
| **N_Caudate-2 L** | 27.16 | 0.002 |
| **N_Caudate-2 R** | 31.97 | 0.001 |
| **N_Caudate-3 L** | 24.82 | 0.002 |
| **N_Caudate-3 R** | 21.56 | 0.002 |
| **N_Caudate-4 L** | 12.51 | 0.011 |
| **N_Caudate-4 R** | 23.86 | 0.002 |
| **N_Caudate-5 L** | 13.67 | 0.009 |
| **N_Caudate-5 R** | 13.79 | 0.008 |
| **N_Caudate-6 L** | 13.68 | 0.009 |
| **N_Caudate-6 R** | 19.63 | 0.003 |
| **N_Caudate-7 L** | 15.85 | 0.006 |
| **N_Caudate-7 R** | 24.02 | 0.002 |
| **N_Pallidum-1 L** | 8.21 | 0.029 |
| **N_Pallidum-1 R** | 18.81 | 0.004 |
| **N_Putamen-2 L** | 17.99 | 0.004 |
| **N_Putamen-2 R** | 22.99 | 0.002 |

| | | |
|---|---|---|
| **N_Putamen-3 L** | 9.19 | 0.022 |
| **N_Putamen-3 R** | 17.50 | 0.004 |
| **N_Thalamus-1 L** | 7.65 | 0.033 |
| **N_Thalamus-1 R** | 6.60 | 0.046 |
| **N_Thalamus-2 L** | 7.78 | 0.032 |
| **N_Thalamus-2 R** | 10.16 | 0.018 |
| **N_Thalamus-5 R** | 15.93 | 0.006 |
| **N_Thalamus-6 L** | 9.55 | 0.020 |
| **N_Thalamus-6 R** | 10.36 | 0.017 |
| **N_Thalamus-7 R** | 12.05 | 0.012 |
| **N_Thalamus-8 L** | 7.19 | 0.037 |
| **N_Thalamus-8 R** | 9.21 | 0.022 |
| **N_Thalamus-9 R** | 7.56 | 0.034 |
| **PPNleft** | 9.40 | 0.022 |
| **PPNright** | 7.03 | 0.039 |
| **S_Anterior_Rostral-1 L** | 31.36 | 0.001 |
| **S_Anterior_Rostral-1 R** | 32.91 | 0.001 |
| **S_Cingulate-1 L** | 25.64 | 0.002 |
| **S_Cingulate-1 R** | 27.23 | 0.002 |
| **S_Cingulate-2 L** | 21.97 | 0.003 |
| **S_Cingulate-2 R** | 33.00 | 0.001 |
| **S_Cingulate-3 L** | 27.10 | 0.002 |
| **S_Cingulate-3 R** | 37.38 | 0.001 |
| **S_Cingulate-4 L** | 32.71 | 0.001 |
| **S_Cingulate-4 R** | 36.27 | 0.001 |
| **S_Cingulate-5 L** | 23.49 | 0.002 |
| **S_Cingulate-5 R** | 59.60 | 0.001 |
| **S_Cingulate-6 L** | 23.80 | 0.002 |
| **S_Cingulate-6 R** | 34.85 | 0.001 |
| **S_Cingulate-7 L** | 17.58 | 0.005 |
| **S_Cingulate-7 R** | 25.82 | 0.002 |
| **S_Inf_Frontal-1 L** | 25.41 | 0.002 |
| **S_Inf_Frontal-1 R** | 21.28 | 0.003 |
| **S_Inf_Frontal-2 L** | 49.24 | 0.001 |
| **S_Inf_Frontal-2 R** | 34.95 | 0.001 |
| **S_Intraparietal-1 L** | 27.71 | 0.002 |
| **S_Intraparietal-1 R** | 23.74 | 0.002 |
| **S_Intraparietal-2 R** | 19.59 | 0.003 |
| **S_Intraparietal-3 R** | 11.64 | 0.013 |
| **S_Olfactory-1 L** | 27.07 | 0.002 |
| **S_Olfactory-1 R** | 21.68 | 0.003 |

| | | |
|---|---|---|
| **S_Orbital-1 L** | 39.14 | 0.001 |
| **S_Orbital-1 R** | 23.04 | 0.002 |
| **S_Orbital-2 L** | 37.41 | 0.001 |
| **S_Orbital-2 R** | 27.47 | 0.002 |
| **S_Postcentral-1 L** | 26.37 | 0.002 |
| **S_Postcentral-1 R** | 63.94 | 0.001 |
| **S_Postcentral-2 L** | 27.69 | 0.002 |
| **S_Postcentral-2 R** | 31.25 | 0.001 |
| **S_Postcentral-3 L** | 29.18 | 0.001 |
| **S_Postcentral-3 R** | 41.04 | 0.001 |
| **S_Precentral-1 L** | 49.60 | 0.001 |
| **S_Precentral-1 R** | 79.33 | 0.001 |
| **S_Precentral-2 L** | 24.35 | 0.002 |
| **S_Precentral-2 R** | 48.08 | 0.001 |
| **S_Precentral-3 L** | 32.27 | 0.001 |
| **S_Precentral-3 R** | 23.63 | 0.002 |
| **S_Precentral-4 L** | 29.31 | 0.001 |
| **S_Precentral-4 R** | 33.25 | 0.001 |
| **S_Precentral-5 L** | 40.16 | 0.001 |
| **S_Precentral-5 R** | 36.03 | 0.001 |
| **S_Precentral-6 L** | 25.84 | 0.002 |
| **S_Precentral-6 R** | 32.10 | 0.001 |
| **S_Rolando-1 L** | 28.53 | 0.001 |
| **S_Rolando-1 R** | 57.70 | 0.001 |
| **S_Rolando-2 L** | 28.82 | 0.002 |
| **S_Rolando-2 R** | 39.37 | 0.001 |
| **S_Rolando-3 L** | 21.87 | 0.003 |
| **S_Rolando-3 R** | 50.13 | 0.001 |
| **S_Rolando-4 L** | 29.85 | 0.001 |
| **S_Rolando-4 R** | 41.99 | 0.001 |
| **S_Sup_Frontal-1 L** | 19.26 | 0.003 |
| **S_Sup_Frontal-1 R** | 14.23 | 0.008 |
| **S_Sup_Frontal-2 L** | 22.07 | 0.002 |
| **S_Sup_Frontal-2 R** | 96.71 | 0.000 |
| **S_Sup_Frontal-3 L** | 28.57 | 0.001 |
| **S_Sup_Frontal-3 R** | 67.52 | 0.001 |
| **S_Sup_Frontal-4 L** | 26.01 | 0.002 |
| **S_Sup_Frontal-4 R** | 26.88 | 0.002 |
| **S_Sup_Frontal-5 L** | 18.96 | 0.004 |
| **S_Sup_Frontal-5 R** | 54.02 | 0.001 |
| **S_Sup_Frontal-6 L** | 26.63 | 0.002 |

| | | |
|---|---|---|
| **S_Sup_Frontal-6 R** | 37.11 | 0.001 |
| **S_Sup_Temporal-1 L** | 36.18 | 0.001 |
| **S_Sup_Temporal-1 R** | 9.78 | 0.019 |
| **S_Sup_Temporal-2 L** | 12.27 | 0.012 |
| **S_Sup_Temporal-2 R** | 30.13 | 0.001 |
| **S_Sup_Temporal-3 L** | 11.62 | 0.013 |
| **S_Sup_Temporal-3 R** | 20.76 | 0.003 |
| **S_Sup_Temporal-4 L** | 10.68 | 0.016 |

**Supplementary Table 4.** Statistically significant regions of interest showing greater theta activity during REM sleep compared to NREM sleep, as determined by LME models. P-values have been corrected for FDR ($p < 0.05$).

| **ROI** | **F-statistic** | **FDR p-value** |
|---|---|---|
| **Cerebelum_10_L** | 25.11 | 0.013 |
| **Cerebelum_10_R** | 14.04 | 0.018 |
| **Cerebelum_3_L** | 24.69 | 0.013 |
| **Cerebelum_3_R** | 13.85 | 0.018 |
| **Cerebelum_4_5_L** | 23.12 | 0.013 |
| **Cerebelum_4_5_R** | 17.56 | 0.014 |
| **Cerebelum_6_L** | 17.91 | 0.013 |
| **Cerebelum_6_R** | 15.56 | 0.016 |
| **Cerebelum_7b_L** | 18.19 | 0.013 |
| **Cerebelum_7b_R** | 17.08 | 0.015 |
| **Cerebelum_8_L** | 15.64 | 0.016 |
| **Cerebelum_8_R** | 18.16 | 0.014 |
| **Cerebelum_9_L** | 13.49 | 0.018 |
| **Cerebelum_9_R** | 15.74 | 0.016 |
| **Cerebelum_Crus1_L** | 14.73 | 0.016 |
| **Cerebelum_Crus1_R** | 14.56 | 0.018 |
| **Cerebelum_Crus2_L** | 13.53 | 0.018 |
| **Cerebelum_Crus2_R** | 16.86 | 0.015 |
| **G_Calcarine-1 L** | 8.33 | 0.041 |
| **G_Calcarine-3 L** | 7.40 | 0.049 |
| **G_Cingulum_Ant-1 L** | 14.59 | 0.016 |
| **G_Cingulum_Ant-1 R** | 13.68 | 0.018 |
| **G_Cingulum_Ant-2 L** | 18.66 | 0.013 |
| **G_Cingulum_Ant-2 R** | 15.00 | 0.016 |
| **G_Cingulum_Mid-1 L** | 7.87 | 0.043 |
| **G_Cingulum_Post-3 R** | 6.97 | 0.050 |
| **G_Cuneus-1 R** | 7.78 | 0.045 |
| **G_Cuneus-2 R** | 7.85 | 0.046 |

| | | |
|---|---|---|
| **G_Frontal_Inf_Orb-1 L** | 12.06 | 0.022 |
| **G_Frontal_Inf_Orb-1 R** | 11.77 | 0.021 |
| **G_Frontal_Inf_Orb-2 L** | 25.95 | 0.013 |
| **G_Frontal_Inf_Orb-2 R** | 11.20 | 0.026 |
| **G_Frontal_Inf_Tri-1 L** | 25.32 | 0.010 |
| **G_Frontal_Inf_Tri-1 R** | 7.71 | 0.046 |
| **G_Frontal_Med_Orb-1 L** | 12.62 | 0.020 |
| **G_Frontal_Med_Orb-1 R** | 8.03 | 0.044 |
| **G_Frontal_Med_Orb-2 L** | 28.27 | 0.013 |
| **G_Frontal_Med_Orb-2 R** | 24.08 | 0.013 |
| **G_Frontal_Mid_Orb-2 L** | 8.60 | 0.040 |
| **G_Frontal_Sup_Medial-1 L** | 15.60 | 0.015 |
| **G_Frontal_Sup_Medial-1 R** | 7.17 | 0.049 |
| **G_Frontal_Sup_Medial-2 L** | 13.53 | 0.012 |
| **G_Frontal_Sup_Medial-2 R** | 8.73 | 0.033 |
| **G_Frontal_Sup_Medial-3 L** | 16.13 | 0.016 |
| **G_Frontal_Sup_Orb-1 L** | 28.76 | 0.012 |
| **G_Frontal_Sup_Orb-1 R** | 14.52 | 0.018 |
| **G_Frontal_Sup-1 L** | 17.19 | 0.012 |
| **G_Frontal_Sup-2 L** | 19.90 | 0.013 |
| **G_Frontal_Sup-2 R** | 14.83 | 0.016 |
| **G_Fusiform-1 L** | 21.42 | 0.013 |
| **G_Fusiform-2 L** | 20.30 | 0.013 |
| **G_Fusiform-2 R** | 9.15 | 0.035 |
| **G_Fusiform-3 L** | 25.35 | 0.013 |
| **G_Fusiform-3 R** | 9.30 | 0.035 |
| **G_Fusiform-4 L** | 22.20 | 0.013 |
| **G_Fusiform-4 R** | 12.18 | 0.021 |
| **G_Fusiform-5 L** | 29.35 | 0.010 |
| **G_Fusiform-5 R** | 12.92 | 0.019 |
| **G_Fusiform-6 L** | 9.91 | 0.030 |
| **G_Fusiform-6 R** | 10.80 | 0.026 |
| **G_Fusiform-7 L** | 10.42 | 0.028 |
| **G_Hippocampus-1 L** | 18.46 | 0.014 |
| **G_Hippocampus-2 L** | 27.78 | 0.012 |
| **G_Insula-anterior-1 L** | 23.16 | 0.013 |
| **G_Insula-anterior-2 L** | 25.70 | 0.013 |
| **G_Insula-anterior-2 R** | 11.16 | 0.025 |
| **G_Insula-anterior-3 L** | 14.93 | 0.016 |
| **G_Insula-anterior-3 R** | 13.60 | 0.019 |
| **G_Insula-anterior-4 L** | 15.25 | 0.016 |

| | | |
|---|---|---|
| **G_Insula-anterior-4 R** | 10.63 | 0.027 |
| **G_Insula-anterior-5 L** | 11.55 | 0.023 |
| **G_Lingual-1 L** | 43.04 | 0.002 |
| **G_Lingual-1 R** | 10.59 | 0.027 |
| **G_Lingual-2 L** | 13.09 | 0.019 |
| **G_Lingual-2 R** | 9.77 | 0.031 |
| **G_Lingual-3 L** | 10.88 | 0.026 |
| **G_Lingual-3 R** | 8.54 | 0.039 |
| **G_Lingual-4 L** | 7.66 | 0.046 |
| **G_Lingual-5 L** | 8.95 | 0.037 |
| **G_Occipital_Inf-1 L** | 11.51 | 0.023 |
| **G_Occipital_Inf-1 R** | 8.91 | 0.037 |
| **G_Occipital_Inf-2 L** | 14.44 | 0.016 |
| **G_Occipital_Inf-2 R** | 12.00 | 0.022 |
| **G_Occipital_Lat-1 L** | 10.70 | 0.027 |
| **G_Occipital_Lat-2 L** | 10.71 | 0.027 |
| **G_Occipital_Lat-3 R** | 7.20 | 0.050 |
| **G_Occipital_Lat-4 L** | 8.42 | 0.041 |
| **G_Occipital_Lat-5 L** | 9.60 | 0.032 |
| **G_Occipital_Lat-5 R** | 8.47 | 0.040 |
| **G_Occipital_Mid-1 R** | 11.67 | 0.025 |
| **G_Occipital_Mid-2 R** | 7.44 | 0.048 |
| **G_Occipital_Mid-3 R** | 7.23 | 0.050 |
| **G_Occipital_Pole-1 L** | 11.77 | 0.023 |
| **G_Occipital_Pole-1 R** | 9.10 | 0.035 |
| **G_Occipital_Sup-1 R** | 7.31 | 0.049 |
| **G_Occipital_Sup-2 R** | 8.80 | 0.038 |
| **G_ParaHippocampal-1 L** | 20.55 | 0.013 |
| **G_ParaHippocampal-1 R** | 8.29 | 0.041 |
| **G_ParaHippocampal-2 L** | 25.97 | 0.013 |
| **G_ParaHippocampal-2 R** | 8.00 | 0.045 |
| **G_ParaHippocampal-3 L** | 23.77 | 0.013 |
| **G_ParaHippocampal-3 R** | 11.97 | 0.022 |
| **G_ParaHippocampal-4 L** | 24.04 | 0.013 |
| **G_ParaHippocampal-4 R** | 10.54 | 0.028 |
| **G_ParaHippocampal-5 L** | 41.72 | 0.003 |
| **G_ParaHippocampal-5 R** | 8.67 | 0.039 |
| **G_Parietal_Sup-4 L** | 8.04 | 0.046 |
| **G_Parietal_Sup-5 L** | 13.07 | 0.024 |
| **G_Precuneus-1 L** | 7.42 | 0.049 |
| **G_Rolandic_Oper-1 L** | 18.06 | 0.013 |

| | | |
|---|---|---|
| **G_Rolandic_Oper-2 L** | 10.99 | 0.019 |
| **G_subcallosal-1 L** | 23.85 | 0.013 |
| **G_subcallosal-1 R** | 22.24 | 0.013 |
| **G_Temporal_Inf-1 L** | 25.24 | 0.013 |
| **G_Temporal_Inf-2 L** | 14.50 | 0.018 |
| **G_Temporal_Inf-3 L** | 14.91 | 0.016 |
| **G_Temporal_Inf-3 R** | 7.87 | 0.045 |
| **G_Temporal_Inf-4 L** | 13.75 | 0.018 |
| **G_Temporal_Inf-4 R** | 7.82 | 0.045 |
| **G_Temporal_Inf-5 L** | 10.75 | 0.027 |
| **G_Temporal_Mid-1 L** | 8.07 | 0.045 |
| **G_Temporal_Mid-2 R** | 7.64 | 0.047 |
| **G_Temporal_Mid-3 L** | 12.50 | 0.023 |
| **G_Temporal_Mid-3 R** | 7.18 | 0.049 |
| **G_Temporal_Pole_Mid-1 L** | 19.80 | 0.013 |
| **G_Temporal_Pole_Mid-2 L** | 15.55 | 0.017 |
| **G_Temporal_Pole_Mid-3 L** | 20.23 | 0.013 |
| **G_Temporal_Pole_Sup-1 L** | 19.83 | 0.013 |
| **G_Temporal_Pole_Sup-2 L** | 19.54 | 0.013 |
| **G_Temporal_Sup-1 L** | 14.57 | 0.018 |
| **G_Temporal_Sup-1 R** | 8.28 | 0.042 |
| **G_Temporal_Sup-4 L** | 7.72 | 0.045 |
| **LGNleft** | 23.70 | 0.013 |
| **medsep** | 13.55 | 0.019 |
| **N_Amygdala-1 L** | 22.81 | 0.013 |
| **N_Caudate-1 L** | 18.10 | 0.013 |
| **N_Caudate-1 R** | 24.11 | 0.013 |
| **N_Caudate-2 L** | 18.30 | 0.013 |
| **N_Caudate-2 R** | 16.40 | 0.015 |
| **N_Caudate-3 L** | 22.36 | 0.013 |
| **N_Caudate-3 R** | 17.20 | 0.015 |
| **N_Caudate-4 L** | 13.89 | 0.018 |
| **N_Caudate-4 R** | 9.90 | 0.030 |
| **N_Caudate-5 L** | 16.66 | 0.015 |
| **N_Caudate-5 R** | 8.68 | 0.039 |
| **N_Caudate-6 L** | 7.56 | 0.046 |
| **N_Pallidum-1 L** | 14.28 | 0.018 |
| **N_Putamen-2 L** | 22.44 | 0.013 |
| **N_Putamen-2 R** | 7.33 | 0.049 |
| **N_Putamen-3 L** | 14.33 | 0.018 |
| **N_Thalamus-1 L** | 12.79 | 0.020 |

| | | |
|---|---|---|
| **N_Thalamus-1 R** | 11.38 | 0.025 |
| **N_Thalamus-2 R** | 10.83 | 0.029 |
| **N_Thalamus-3 L** | 11.36 | 0.025 |
| **N_Thalamus-3 R** | 10.40 | 0.028 |
| **N_Thalamus-4 R** | 8.20 | 0.045 |
| **N_Thalamus-5 R** | 8.31 | 0.045 |
| **N_Thalamus-9 L** | 7.56 | 0.049 |
| **N_Thalamus-9 R** | 8.48 | 0.041 |
| **PPNleft** | 19.32 | 0.013 |
| **PPNright** | 14.23 | 0.018 |
| **S_Anterior_Rostral-1 L** | 19.08 | 0.013 |
| **S_Anterior_Rostral-1 R** | 15.67 | 0.016 |
| **S_Cingulate-1 L** | 16.65 | 0.015 |
| **S_Cingulate-1 R** | 9.81 | 0.035 |
| **S_Inf_Frontal-2 L** | 7.94 | 0.045 |
| **S_Intraoccipital-1 L** | 9.99 | 0.032 |
| **S_Intraoccipital-1 R** | 15.30 | 0.019 |
| **S_Olfactory-1 L** | 33.90 | 0.012 |
| **S_Olfactory-1 R** | 20.79 | 0.013 |
| **S_Orbital-1 L** | 20.94 | 0.013 |
| **S_Orbital-1 R** | 11.14 | 0.026 |
| **S_Orbital-2 L** | 14.89 | 0.017 |
| **S_Orbital-2 R** | 12.74 | 0.019 |
| **S_Parietooccipital-1 L** | 13.00 | 0.019 |
| **S_Parietooccipital-2 L** | 9.31 | 0.034 |
| **S_Parietooccipital-2 R** | 7.71 | 0.046 |
| **S_Parietooccipital-4 L** | 12.55 | 0.020 |
| **S_Parietooccipital-5 R** | 9.74 | 0.031 |
| **S_Postcentral-1 L** | 10.67 | 0.021 |
| **S_Rolando-2 L** | 7.83 | 0.050 |
| **S_Sup_Frontal-1 L** | 12.41 | 0.020 |
| **S_Sup_Frontal-1 R** | 11.11 | 0.025 |
| **S_Sup_Frontal-2 L** | 13.62 | 0.018 |
| **S_Sup_Frontal-3 L** | 12.66 | 0.019 |
| **S_Sup_Frontal-3 R** | 7.49 | 0.049 |
| **S_Sup_Frontal-4 L** | 6.94 | 0.035 |
| **S_Sup_Temporal-1 L** | 19.86 | 0.013 |
| **S_Sup_Temporal-3 L** | 17.52 | 0.016 |
| **Vermis_1_2** | 18.17 | 0.013 |
| **Vermis_10** | 13.34 | 0.019 |
| **Vermis_3** | 13.44 | 0.018 |

| | | |
|---|---|---|
| **Vermis_4_5** | 9.94 | 0.029 |
| **Vermis_6** | 13.22 | 0.019 |
| **Vermis_7** | 16.47 | 0.015 |
| **Vermis_8** | 17.42 | 0.015 |
| **Vermis_9** | 16.37 | 0.015 |

**Supplementary Table 5.** Statistically significant regions of interest showing greater theta activity during tonic REM sleep compared to phasic REM sleep, as determined by LME models. P-values have been corrected for FDR ($p < 0.05$).

| ROI | F-statistic | FDR p-value |
|---|---|---|
| **PPNleft** | 19.321 | 0.013 |
| **PPNright** | 14.227 | 0.018 |
| **Cerebelum_10_L** | 25.111 | 0.013 |
| **Cerebelum_10_R** | 14.043 | 0.018 |
| **Cerebelum_3_L** | 24.685 | 0.013 |
| **Cerebelum_3_R** | 13.852 | 0.018 |
| **Cerebelum_4_5_L** | 23.118 | 0.013 |
| **Cerebelum_4_5_R** | 17.558 | 0.014 |
| **Cerebelum_6_L** | 17.914 | 0.013 |
| **Cerebelum_6_R** | 15.562 | 0.016 |
| **Cerebelum_7b_L** | 18.187 | 0.013 |
| **Cerebelum_7b_R** | 17.078 | 0.015 |
| **Cerebelum_8_L** | 15.637 | 0.016 |
| **Cerebelum_8_R** | 18.160 | 0.014 |
| **Cerebelum_9_L** | 13.495 | 0.018 |
| **Cerebelum_9_R** | 15.740 | 0.016 |
| **Cerebelum_Crus1_L** | 14.727 | 0.016 |
| **Cerebelum_Crus1_R** | 14.556 | 0.018 |
| **Cerebelum_Crus2_L** | 13.531 | 0.018 |
| **Cerebelum_Crus2_R** | 16.864 | 0.015 |
| **G_Calcarine-1 L** | 8.331 | 0.041 |
| **G_Calcarine-3 L** | 7.403 | 0.049 |
| **G_Cingulum_Ant-1 L** | 14.586 | 0.016 |
| **G_Cingulum_Ant-1 R** | 13.678 | 0.018 |
| **G_Cingulum_Ant-2 L** | 18.663 | 0.013 |
| **G_Cingulum_Ant-2 R** | 14.996 | 0.016 |
| **G_Cingulum_Mid-1 L** | 7.869 | 0.043 |
| **G_Cingulum_Post-3 R** | 6.971 | 0.050 |
| **G_Cuneus-1 R** | 7.781 | 0.045 |
| **G_Cuneus-2 R** | 7.852 | 0.046 |

| | | |
|---|---|---|
| **G_Frontal_Inf_Orb-1 L** | 12.058 | 0.022 |
| **G_Frontal_Inf_Orb-1 R** | 11.768 | 0.021 |
| **G_Frontal_Inf_Orb-2 L** | 25.951 | 0.013 |
| **G_Frontal_Inf_Orb-2 R** | 11.197 | 0.026 |
| **G_Frontal_Inf_Tri-1 L** | 25.320 | 0.010 |
| **G_Frontal_Inf_Tri-1 R** | 7.712 | 0.046 |
| **G_Frontal_Med_Orb-1 L** | 12.619 | 0.020 |
| **G_Frontal_Med_Orb-1 R** | 8.032 | 0.044 |
| **G_Frontal_Med_Orb-2 L** | 28.267 | 0.013 |
| **G_Frontal_Med_Orb-2 R** | 24.078 | 0.013 |
| **G_Frontal_Mid_Orb-2 L** | 8.597 | 0.040 |
| **G_Frontal_Sup-1 L** | 17.192 | 0.012 |
| **G_Frontal_Sup-2 L** | 19.904 | 0.013 |
| **G_Frontal_Sup-2 R** | 14.826 | 0.016 |
| **G_Frontal_Sup_Medial-1 L** | 15.601 | 0.015 |
| **G_Frontal_Sup_Medial-1 R** | 7.174 | 0.049 |
| **G_Frontal_Sup_Medial-2 L** | 13.526 | 0.012 |
| **G_Frontal_Sup_Medial-2 R** | 8.732 | 0.033 |
| **G_Frontal_Sup_Medial-3 L** | 16.129 | 0.016 |
| **G_Frontal_Sup_Orb-1 L** | 28.759 | 0.012 |
| **G_Frontal_Sup_Orb-1 R** | 14.516 | 0.018 |
| **G_Fusiform-1 L** | 21.417 | 0.013 |
| **G_Fusiform-2 L** | 20.300 | 0.013 |
| **G_Fusiform-2 R** | 9.153 | 0.035 |
| **G_Fusiform-3 L** | 25.348 | 0.013 |
| **G_Fusiform-3 R** | 9.297 | 0.035 |
| **G_Fusiform-4 L** | 22.198 | 0.013 |
| **G_Fusiform-4 R** | 12.184 | 0.021 |
| **G_Fusiform-5 L** | 29.350 | 0.010 |
| **G_Fusiform-5 R** | 12.923 | 0.019 |
| **G_Fusiform-6 L** | 9.914 | 0.030 |
| **G_Fusiform-6 R** | 10.802 | 0.026 |
| **G_Fusiform-7 L** | 10.422 | 0.028 |
| **G_Hippocampus-1 L** | 18.464 | 0.014 |
| **G_Hippocampus-2 L** | 27.775 | 0.012 |
| **G_Insula-anterior-1 L** | 23.162 | 0.013 |
| **G_Insula-anterior-2 L** | 25.704 | 0.013 |
| **G_Insula-anterior-2 R** | 11.162 | 0.025 |
| **G_Insula-anterior-3 L** | 14.930 | 0.016 |
| **G_Insula-anterior-3 R** | 13.598 | 0.019 |
| **G_Insula-anterior-4 L** | 15.250 | 0.016 |

| | | |
|---|---|---|
| **G_Insula-anterior-4 R** | 10.635 | 0.027 |
| **G_Insula-anterior-5 L** | 11.545 | 0.023 |
| **G_Lingual-1 L** | 43.040 | 0.002 |
| **G_Lingual-1 R** | 10.587 | 0.027 |
| **G_Lingual-2 L** | 13.087 | 0.019 |
| **G_Lingual-2 R** | 9.775 | 0.031 |
| **G_Lingual-3 L** | 10.881 | 0.026 |
| **G_Lingual-3 R** | 8.535 | 0.039 |
| **G_Lingual-4 L** | 7.664 | 0.046 |
| **G_Lingual-5 L** | 8.953 | 0.037 |
| **G_Occipital_Inf-1 L** | 11.510 | 0.023 |
| **G_Occipital_Inf-1 R** | 8.906 | 0.037 |
| **G_Occipital_Inf-2 L** | 14.441 | 0.016 |
| **G_Occipital_Inf-2 R** | 12.001 | 0.022 |
| **G_Occipital_Lat-1 L** | 10.697 | 0.027 |
| **G_Occipital_Lat-2 L** | 10.713 | 0.027 |
| **G_Occipital_Lat-3 R** | 7.201 | 0.050 |
| **G_Occipital_Lat-4 L** | 8.415 | 0.041 |
| **G_Occipital_Lat-5 L** | 9.599 | 0.032 |
| **G_Occipital_Lat-5 R** | 8.475 | 0.040 |
| **G_Occipital_Mid-1 R** | 11.667 | 0.025 |
| **G_Occipital_Mid-2 R** | 7.439 | 0.048 |
| **G_Occipital_Mid-3 R** | 7.230 | 0.050 |
| **G_Occipital_Pole-1 L** | 11.774 | 0.023 |
| **G_Occipital_Pole-1 R** | 9.101 | 0.035 |
| **G_Occipital_Sup-1 R** | 7.305 | 0.049 |
| **G_Occipital_Sup-2 R** | 8.803 | 0.038 |
| **G_ParaHippocampal-1 L** | 20.553 | 0.013 |
| **G_ParaHippocampal-1 R** | 8.286 | 0.041 |
| **G_ParaHippocampal-2 L** | 25.975 | 0.013 |
| **G_ParaHippocampal-2 R** | 7.997 | 0.045 |
| **G_ParaHippocampal-3 L** | 23.775 | 0.013 |
| **G_ParaHippocampal-3 R** | 11.965 | 0.022 |
| **G_ParaHippocampal-4 L** | 24.036 | 0.013 |
| **G_ParaHippocampal-4 R** | 10.537 | 0.028 |
| **G_ParaHippocampal-5 L** | 41.720 | 0.003 |
| **G_ParaHippocampal-5 R** | 8.672 | 0.039 |
| **G_Parietal_Sup-4 L** | 8.040 | 0.046 |
| **G_Parietal_Sup-5 L** | 13.068 | 0.024 |
| **G_Precuneus-1 L** | 7.418 | 0.049 |
| **G_Rolandic_Oper-1 L** | 18.057 | 0.013 |

| | | |
|---|---|---|
| **G_Rolandic_Oper-2 L** | 10.988 | 0.019 |
| **G_subcallosal-1 L** | 23.854 | 0.013 |
| **G_subcallosal-1 R** | 22.236 | 0.013 |
| **G_Temporal_Inf-1 L** | 25.242 | 0.013 |
| **G_Temporal_Inf-2 L** | 14.499 | 0.018 |
| **G_Temporal_Inf-3 L** | 14.915 | 0.016 |
| **G_Temporal_Inf-3 R** | 7.870 | 0.045 |
| **G_Temporal_Inf-4 L** | 13.755 | 0.018 |
| **G_Temporal_Inf-4 R** | 7.816 | 0.045 |
| **G_Temporal_Inf-5 L** | 10.751 | 0.027 |
| **G_Temporal_Mid-1 L** | 8.069 | 0.045 |
| **G_Temporal_Mid-2 R** | 7.645 | 0.047 |
| **G_Temporal_Mid-3 L** | 12.495 | 0.023 |
| **G_Temporal_Mid-3 R** | 7.184 | 0.049 |
| **G_Temporal_Pole_Mid-1 L** | 19.798 | 0.013 |
| **G_Temporal_Pole_Mid-2 L** | 15.555 | 0.017 |
| **G_Temporal_Pole_Mid-3 L** | 20.230 | 0.013 |
| **G_Temporal_Pole_Sup-1 L** | 19.831 | 0.013 |
| **G_Temporal_Pole_Sup-2 L** | 19.535 | 0.013 |
| **G_Temporal_Sup-1 L** | 14.573 | 0.018 |
| **G_Temporal_Sup-1 R** | 8.280 | 0.042 |
| **G_Temporal_Sup-4 L** | 7.719 | 0.045 |
| **medsep** | 13.545 | 0.019 |
| **N_Amygdala-1 L** | 22.808 | 0.013 |
| **N_Caudate-1 L** | 18.101 | 0.013 |
| **N_Caudate-1 R** | 24.105 | 0.013 |
| **N_Caudate-2 L** | 18.300 | 0.013 |
| **N_Caudate-2 R** | 16.403 | 0.015 |
| **N_Caudate-3 L** | 22.356 | 0.013 |
| **N_Caudate-3 R** | 17.196 | 0.015 |
| **N_Caudate-4 L** | 13.890 | 0.018 |
| **N_Caudate-4 R** | 9.901 | 0.030 |
| **N_Caudate-5 L** | 16.662 | 0.015 |
| **N_Caudate-5 R** | 8.682 | 0.039 |
| **N_Caudate-6 L** | 7.564 | 0.046 |
| **N_Pallidum-1 L** | 14.277 | 0.018 |
| **N_Putamen-2 L** | 22.436 | 0.013 |
| **N_Putamen-2 R** | 7.330 | 0.049 |
| **N_Putamen-3 L** | 14.334 | 0.018 |
| **N_Thalamus-1 L** | 12.792 | 0.020 |
| **N_Thalamus-1 R** | 11.375 | 0.025 |

| | | |
|---|---|---|
| **N_Thalamus-2 R** | 10.830 | 0.029 |
| **N_Thalamus-3 L** | 11.358 | 0.025 |
| **N_Thalamus-3 R** | 10.395 | 0.028 |
| **N_Thalamus-4 R** | 8.202 | 0.045 |
| **N_Thalamus-5 R** | 8.308 | 0.045 |
| **N_Thalamus-9 L** | 7.565 | 0.049 |
| **N_Thalamus-9 R** | 8.485 | 0.041 |
| **S_Anterior_Rostral-1 L** | 19.079 | 0.013 |
| **S_Anterior_Rostral-1 R** | 15.670 | 0.016 |
| **S_Cingulate-1 L** | 16.647 | 0.015 |
| **S_Cingulate-1 R** | 9.814 | 0.035 |
| **S_Inf_Frontal-2 L** | 7.936 | 0.045 |
| **S_Intraoccipital-1 L** | 9.989 | 0.032 |
| **S_Intraoccipital-1 R** | 15.304 | 0.019 |
| **S_Olfactory-1 L** | 33.901 | 0.012 |
| **S_Olfactory-1 R** | 20.792 | 0.013 |
| **S_Orbital-1 L** | 20.941 | 0.013 |
| **S_Orbital-1 R** | 11.138 | 0.026 |
| **S_Orbital-2 L** | 14.888 | 0.017 |
| **S_Orbital-2 R** | 12.742 | 0.019 |
| **S_Parietooccipital-1 L** | 12.997 | 0.019 |
| **S_Parietooccipital-2 L** | 9.306 | 0.034 |
| **S_Parietooccipital-2 R** | 7.707 | 0.046 |
| **S_Parietooccipital-4 L** | 12.546 | 0.020 |
| **S_Parietooccipital-5 R** | 9.739 | 0.031 |
| **S_Postcentral-1 L** | 10.670 | 0.021 |
| **S_Rolando-2 L** | 7.830 | 0.050 |
| **S_Sup_Frontal-1 L** | 12.407 | 0.020 |
| **S_Sup_Frontal-1 R** | 11.107 | 0.025 |
| **S_Sup_Frontal-2 L** | 13.625 | 0.018 |
| **S_Sup_Frontal-3 L** | 12.660 | 0.019 |
| **S_Sup_Frontal-3 R** | 7.488 | 0.049 |
| **S_Sup_Frontal-4 L** | 6.944 | 0.035 |
| **S_Sup_Temporal-1 L** | 19.861 | 0.013 |
| **S_Sup_Temporal-3 L** | 17.516 | 0.016 |
| **Vermis_10** | 13.340 | 0.019 |
| **Vermis_1_2** | 18.166 | 0.013 |
| **Vermis_3** | 13.437 | 0.018 |
| **Vermis_4_5** | 9.938 | 0.029 |
| **Vermis_6** | 13.221 | 0.019 |
| **Vermis_7** | 16.472 | 0.015 |

| | | |
|---|---|---|
| **Vermis_8** | 17.418 | 0.015 |
| **Vermis_9** | 16.371 | 0.015 |
| **LGNleft** | 23.700 | 0.013 |
